# Isolation and proteomic analysis of intracellular vesicles from the potato late blight pathogen *Phytophthora infestans*

**DOI:** 10.1101/2025.09.30.679036

**Authors:** Jasmine Pham, Stephen C Whisson, Charlotte H Hurst, Sean Chapman, Paul RJ Birch

**Affiliations:** Division of Molecular Cell and Developmental Biology, School of Life Sciences, University of Dundee, DD2 5DA, UK; Division of Plant Sciences, University of Dundee, at The James Hutton Institute, Errol Rd, Invergowrie, Dundee, DD2 5DA, UK; Department of Cell and Molecular Sciences, The James Hutton Institute, Invergowrie, Dundee, DD2 5DA; Medical Research Council Protein Phosphorylation and Ubiquitylation Unit, School of Life Sciences, University of Dundee, DD2 5DA, UK

**Keywords:** *Phytophthora infestans*, effector proteins, protein secretion, vesicles, membrane proteins, density gradient ultracentrifugation, iodixanol, DIA mass spectrometry

## Abstract

The oomycete *Phytophthora infestans*, a filamentous plant pathogen belonging to the kingdom Stramenopila, is the causal agent of potato late blight resulting in annual crop losses amounting to billions of dollars worldwide. Key to the success of this pathogen are the effector proteins it secretes during infection, whose functions include breaching the plant cell wall and suppression/evasion of plant immune responses. Currently, little is known regarding the intracellular trafficking of effectors enroute to secretion. In this study, we developed a robust density gradient ultracentrifugation method to isolate intracellular vesicles from *P. infestans* and to separate diverse vesicle populations based on buoyancy for the purpose of identifying vesicle-associated proteins by mass spectrometry. Gene Ontology Enrichment Analysis of proteins identified in buoyant fractions revealed enrichment for membrane-associated proteins and proteins involved in vesicle trafficking. Buoyant fractions were also enriched in RXLR class effectors, carbohydrate-active enzymes, and secretory proteins possessing N-terminal signal peptides, all representing potential vesicle cargo. In addition, previously identified *P. infestans* extracellular vesicle markers were also present. Unravelling how effector proteins are trafficked for secretion during infection is a critical step in developing robust strategies for combating potato late blight disease; the proteomics dataset and method presented here are valuable resources from which potential biomarkers for *P. infestans* vesicles can be identified for future studies towards this end.

## Introduction

*Phytophthora infestans* is a major pathogen of potato and tomato, causing annual losses amounting to more than €1 billion within the European Union alone ^1^. *P. infestans* is notoriously difficult to control in the field, with current control methods based on costly fungicides, some with unknown modes of action. Attempts to breed crops with long-lasting resistance have generally been unsuccessful ^1,2^. Therefore, unravelling how *P. infestans* invades host plant cells is a critical step in developing robust strategies for combating this important plant pathogen.

*P. infestans* is an oomycete belonging to the kingdom Stramenopila, and although resembling fungi in appearance, the oomycetes are distinguished by their coenocytic hyphae lacking septa, and the presence of cellulose rather than chitin in their cell walls ^3^. During infection, *P. infestans* develops haustoria, finger-like projections that form intimate interactions with the plant cell membrane. It is at this pathogen-plant interface that *P. infestans* secretes and delivers a range of proteins, small molecules and RNAs. Their functions range from assisting evasion of plant immunity, to physically breaching plant tissue and nutrient uptake (reviewed in Boevink *et al.* ^4^).

Amongst the proteins secreted during infection by *P. infestans,* effector proteins have been the focus of intense research and can be divided into two broad classes: apoplastic (extracellular) and cytoplasmic (intracellular). Apoplastic effectors, which act within the apoplastic space outside the plant cell, often function as hydrolytic enzymes/cell wall degrading enzymes, such as pectin esterases ^5^. Other apoplastic effector functions include nutrient scavenging, as is the case for elicitins which are reported to facilitate sterol and lipid acquisition ^6^; various proteases (e.g., aspartic, cysteine and metalloproteases) (reviewed in Saraiva *et al.* ^7^); and inhibitors of plant hydrolases and proteases ^8–10^ secreted in defence against *P. infestans*.

In contrast, cytoplasmic effectors are translocated into the plant cell. The most well studied and abundant in *P. infestans* are the RxLR effectors, with 563 encoded within the genome ^11^. RxLR effectors are characterised by the presence of an Arg-x-Leu-Arg motif (with ‘x’ denoting any amino acid) downstream of a signal peptide near the N terminus. RxLR effector protein function arises from their variable C-termini and accounts for their wide-ranging functions, which include interference/modulation of host signalling, protein regulation, transcription, RNA trafficking and processing, and cellular trafficking ^12^.

Given the importance of proteins secreted by *P. infestans* in modulating the outcome of infection and disease, much attention has focussed on the fate and function of these proteins once secreted into the host plant. However, the intracellular trafficking pathways enroute to extracellular secretion remain understudied. The ability to track the intracellular pathways of effectors would aid in developing novel targeted control methods to disrupt effector secretion and inhibit infection. Achieving this will require the identification of markers for intracellular compartments and vesicles through, and by, which these proteins are trafficked.

A bottleneck for studying protein secretion in *P. infestans,* and indeed other filamentous plant pathogens, is the lack of well-characterised vesicle markers such as those found in mammalian systems ^13,14^. Notably, tetraspanins, a group of well conserved multi-pass transmembrane proteins routinely used as markers for vesicles in mammalian research ^13^, have only recently been identified in *Phytophthora sojae* extracellular vesicles ^15^. In *P. infestans*, two transmembrane MARVEL domain proteins have emerged as potential markers for extracellular vesicles (EVs) associated with RXLR effectors ^16^.

Despite the progress in identifying EV-associated proteins, little research has been carried out to identify intracellular vesicle proteins. Here, we present a robust method for isolating intracellular vesicles from *P. infestans* to identify potential marker and cargo proteins by mass spectrometry. By combining differential centrifugation and cushioned ultracentrifugation with bottom-loaded gradient ultracentrifugation, we isolated intracellular *P. infestans* vesicles based on buoyancy. Isolated proteins were then identified by data-independent acquisition (DIA) mass spectrometry to generate a robust dataset of intracellular vesicle-associated proteins.

This dataset represents the first community resource to identify markers for both intracellular compartment vesicles and cargo proteins from *Phytophthora infestans* for future studies. The robust method presented here can also be adapted in future studies to investigate specific intracellular vesicle populations in more detail. This resource will enable dissection of the intracellular trafficking pathways of effector proteins, with a view to identifying novel methods for controlling this devastating potato pathogen.

## Materials and methods

### Phytophthora infestans in vitro culture

For culturing *in vitro* on solid media, *P. infestans* strain 3928A was cultured on rye agar plates supplemented with 100 µg/mL ampicillin and 10 µg/mL pimaricin at 19°C in the dark for 11 to 14 days for sporulation ^17^. For transformants, plates were additionally supplemented with 10 µg/mL G418.

For culturing in liquid media, sporangia from *P. infestans* were collected by flooding plates with amended Lima bean broth (ALB) ^17^ supplemented with 100 µg/mL ampicillin (and additionally 2.5 µg/mL G418 for transformants) and scraping the plates with a spreader. The media was then filtered through a 70 µM cell sieve (Corning) or a filter unit fitted with 70 µM nylon mesh to remove mycelial fragments.

For confirmation of expression and secretion of fluorescently tagged proteins in transformants, sporangia were collected from two 9 cm diameter Petri dish cultures into a total of 10 mL ALB as described above. The filtered sporangia were incubated in the dark for three days at 19°C. Mycelia were then transferred to 1 mL clarified ALB (centrifuged at 10,000 × g for 2 h to remove particulate matter) and incubated in the dark for a further 24 h, after which the mycelia and condition media (CM) were collected. Mycelia were dried on tissue to remove excess media, and proteins in the CM were chloroform-methanol precipitated. 2 × SDS loading buffer was added directly to the mycelial sample (60 µl) and to the chloroform-precipitated protein pellet (40 µl) and processed for western blotting as described below.

For culturing for intracellular vesicle isolation by cushioned gradient ultracentrifugation, sporangia from 42 × 15 cm diameter Petri dish cultures were collected and used to inoculate 6 × 100 mL of clarified ALB as described above. Cultures were grown at 19°C in the dark for five days and mycelia collected and processed for intracellular vesicle isolation as detailed below.

### Expression of fluorescently tagged effectors in *P. infestans*

A dual expression vector for simultaneous expression of both an mCherry tagged protein with an mCitrine tagged protein was generated by modification of the vector pmCherryN ^18^. mCitrine was amplified by PCR from vector pmCitrine_N1 (Addgene, plasmid #92420) to introduce the restriction endonuclease sites ClaI-SpeI-AsiSI flanking the N- and C-termini of mCitrine, respectively. The PCR product was cloned into the vector pTOR (NCBI GenBank accession EU257520) between the ClaI and EcoRI restriction sites. The pTOR vector contains a Ham34 promoter (HamP) and a Ham34 terminator (HamT) up- and downstream of the multiple cloning site (respectively) for expression in *Phytophthora.* The insert with the upstream HamP and downstream HamT was cloned with primers designed to introduce flanking SacI sites; this SacI-HamP-ClaI-SpeI-AsiSI-mCitrine-XbaI-FseI-EcoRI-HamT-SacI expression cassette was inserted into the SacI site of pmCherryN to generate the dual expression vector pDual_mCherryN_mCitrine (Supplementary Figure S1).

The RxLR effector PITG_04314 was cloned between the AgeI/PacI restriction sites upstream of the mCherry gene, and the pectinesterase PITG_01029 was cloned between the ClaI/AsiSI restriction sites upstream of mCitrine. All primers used are listed in Supplementary Table 1.

The dual vector containing PITG_04314 and PITG_01029 was transformed into *P. infestans* protoplasts using a polyethylene glycol (PEG)-CaCl_2_-lipofectin mediated method ^17,19^.

### Plant growth conditions

*Nicotiana benthamiana* was grown in commercially available compost under glasshouse conditions of 16/8 h light/dark cycle with a maximum daytime temperature of 26 °C and night-time temperature of 22 °C. Plants were used at 4-5 weeks old.

### P. infestans inoculation of N. benthamiana

*P. infestans* sporangia were harvested in sterile distilled water from 11 - 14 days old cultures, passed through a 70 µm cell sieve (Corning), and the concentration adjusted to 5 × 10^5^ sporangia per mL for use as inoculum. The underside of detached *N. benthamiana* leaves were nicked with a scalpel blade and a 10 µL drop of inoculum deposited at this site. The leaves were incubated at 19°C in a humid box and imaged by confocal microscopy at 5 days post inoculation (dpi).

### Confocal microscopy

*N. benthamiana* leaves infected with transgenic *P. infestans* co-expressing PITG_04314-mCherry and PITG_01029-mCitrine were mounted on slides and imaged using a Nikon A1R confocal microscope with a 40 × water immersion lens. mCherry was imaged using 561 nm excitation and emissions collected between 570 and 620 nm. mCitrine was imaged with 488 nm excitation and emissions collected between 500 and 530 nm.

### Intracellular vesicle isolation

Mycelia grown in 100 mL liquid culture (as described above) was collected using a filter unit fitted with 70 µm nylon mesh and rinsed with cold isolation buffer adapted from Heard *et al*. ^20^ (150 mM Na-HEPES, 10 mM M EDTA, 10 mM EGTA, 17.5 % (w/v) sucrose, 7.5 mM KCl, 1 × protease inhibitor (Roche)). Mycelia were then ground with a pestle and mortar with 11 mL isolation buffer and filtered through a 40 µm cell sieve to remove large debris. The extract was pre-cleared by differential centrifugation at 2,000 × g for 10 minutes, followed by centrifugation at 10,000 × g for 30 minutes before cushioned ultracentrifugation. All steps were performed on ice or at 4°C.

### Cushioned bottom-loaded density gradient ultracentrifugation

10 mL of pre-cleared extract was overlaid onto a 2 mL 60 % Iodixanol/OptiPrep (STEMCELL Technologies) cushion in a 14 × 89 mm polycarbonate open top tube (Beckman Coulter) and centrifuged at 100,000 × g for 2 h in a Beckman Coulter Optima L-80 XP ultracentrifuge with a swinging bucket rotor (SW41 Ti). After centrifugation, all but 1 mL of extract above the cushion was discarded. The remaining 1 mL of extract was combined with the 2 mL iodixanol cushion and transferred to the bottom of a new ultracentrifuge tube. The sample was overlaid sequentially with 1.3 mL each of 35 %, 30 %, 25 %, 20 %, 17.5 %, 15 % and 10 % iodixanol to form a discontinuous gradient and centrifuged at 120,000 × g for 20h without braking during deceleration. Starting from the top of the tube, eight 1.5 mL fractions (F1-F8) were collected from the gradient for further analysis and processing. All steps were performed on ice or at 4°C. OD 340 nm readings from fractions were measured using a Nanodrop1 spectrophotometer and densities calculated using a standard curve of iodixanol dilutions of known densities.

### Triton X-100 treatment

1% Triton X-100 was added to the 1 mL extract combined with the 2 mL iodixanol cushion after cushioned ultracentrifugation (described above) and incubated on ice for 30 minutes before overlaying with iodixanol as above for gradient ultracentrifugation. A control sample which was incubated on ice for 30 min without the addition of 1% Triton X-100 was run in parallel.

### Deglycosylation assay

Mycelia and culture medium of the dual transformant grown in 75 mL of clarified ALB for five days was collected. Mycelia were ground in 5 mL of TEN buffer (25 mM Tris-HCl, pH 7.5, 1 mM EDTA, 150 mM NaCl) and centrifuged for 10 minutes at 3,500 × g to remove cellular debris. 5 ml mycelial extract and 5 ml of the culture medium were precipitated by chloroform-methanol extraction and the protein pellet resuspended in 500 µl sterile distilled water. 40 µl of each sample was treated with Protein Deglycosylation Mix II (NEB) under denaturing conditions following the manufacturer’s instructions. 2 × LDS loading buffer was added to each sample and analysed by western blot as described below.

### Western blotting

Protein samples in 2 × SDS loading buffer were incubated at 95°C for 10 min; protein samples in 2 × LDS loading buffer were incubated at 70°C for 10 minutes. Samples were separated on 12 % acrylamide tris-glycine gels and transferred onto 0.45 µm nitrocellulose membranes (Amersham). Total proteins on membranes were visualised using Revert 700 Total Protein Stain Kit (LI-COR) following manufacturer’s instructions. Membranes were blocked in 4% milk in PBS and probed with rat anti-mRFP (Chromotek) and mouse anti-GFP (Roche) antibodies for PITG_04314-mCherry and PITG_01029-mCitrine, respectively. Membranes were then probed with IRDye® 800CW Goat anti-Mouse IgG and IRDye® 800CW Goat anti-Rat IgG secondary antibodies (LI-COR) and imaged with the LI-COR Odyssey CLx Near-Infrared Fluorescence Imaging System. Images were analysed/quantified using Image Studio Lite version 5.2.5 (LI-COR).

### Transmission electron microscopy of intracellular vesicles in gradient fractions

Samples were fixed with 2 % formaldehyde and 1% glutaraldehyde onto carbon coated formvar on 300 mesh nickel grids (Agar Scientific), stained with 4% uranyl oxylate (pH 7) and embedded with 0.15 % methyl cellulose and 0.4 % uranyl acetate. Grids were imaged with a Jeol JEM 1400 transmission electron microscope.

### Processing of gradient fractions for mass spectrometry

Equivalent fractions from six gradients performed simultaneously for fractions F2, F3, F4 and F6 were pooled to form a single replicate experiment; a total of four replicate experiments performed on four separate occasions were processed. Proteins were chloroform-methanol precipitated, and protein pellets resuspended in 200 µl 1% sodium deoxycholate (Sigma-Aldrich), 50 mM Tris-HCl, pH 8. Proteins were reduced with 5 mM DTT and alkylated with 18.75 mM Iodoacetamide and digested with 2 µg Trypsin/Lys-C mix (Promega) overnight at 37°C in the presence of 1% sodium deoxycholate. The samples were then further digested with 1 µg Trypsin Platinum (Promega) for 3 h. Detergent was removed from the peptides by ethyl acetate extraction ^21^ followed by processing with HiPPR™ Detergent Removal Spin Column Kit (Thermo Fisher Scientific). Mass spectrometry of the samples was performed by the FingerPrints Proteomics Facility at the University of Dundee.

### DIA LC-MS/MS

Analysis of peptide readout was performed on a Q Exactive HF, Mass Spectrometer (Thermo Scientific) coupled to a Dionex Ultimate 3000 RS (Thermo Scientific). LC buffers used are the following: buffer A (0.1% formic acid in Milli-Q water (v/v), buffer B (80% acetonitrile and 0.1% formic acid in Milli-Q water (v/v), loading buffer (0.1% TF in Milli-Q). Aliquots of 5 or 10 µL desalted peptides were loaded at 10 μL/min onto a trap column (100 μm × 2 cm, PepMap nanoViper C18 column, 5 μm, 100 Å, Thermo Scientific) equilibrated in 0.1% TFA for 15min. The trapping column was washed for 6 min at the same flow rate with 0.1% TFA and then switched in-line with µPAC Neo, 110 cm, nanoLC column, equilibrated at a flow rate of 300nl/min for 30 min. The peptides were eluted from the column at a constant flow rate of 300 nl/min with a linear gradient from 1% buffer B to 7% buffer B in 6 min, from 7% B to 26% buffer B in 117 min, from 26% buffer B to 36% buffer B within 23 min and then from 46 % buffer B to 98% buffer B in 10 min. The column was then washed with 98% buffer B for 18 min. Two blanks were run between each sample to reduce carry-over. The column was kept at a constant temperature of 50°C.

Q-exactive HF was operated in positive ionization mode using an easy spray source. The source voltage was set to 2.2 Kv and the capillary temperature was 250°C. Data were acquired in Data Independent Acquisition Mode as previously described ^22^, with little modification. A scan cycle comprised a full MS scan (*m/z* range from 345-155), resolution was set to 60,000, AGC target 3 × 106, maximum injection time 200 ms. MS survey scans were followed by DIA scans of dynamic window widths with an overlap of 0.5 Th. DIA spectra were recorded at a resolution of 30.000 at 200 *m/z* using an automatic gain control target of 3 × 106, a maximum injection time of 55 ms and a first fixed mass of 200 *m/z*. Normalised collision energy was set to 25 % with a default charge state set at 3. Data for both MS scan and MS/MS DIA scan events were acquired in profile mode.

### Data Analysis

The Thermo raw files were analysed using Spectronaut (version 17.4.230317.55965, Biognosys, AG). The directDIA workflow, using the default settings (BGS Factory Settings) with the following modifications: decoy generation set to inverse; Protein LFQ Method was set to QUANT 2.0 (SN Standard) and Precursor Filtering set to Identified (Qvalue); Cross-Run Normalization was unchecked; Precursor Qvalue Cutoff and Protein Qvalue Cutoff (Experimental) set to 0.01; Precursor PEP Cutoff set to 0.01, Protein Qvalue Cutoff (Run) set to 0.01 and Protein PEP Cutoff set to 0.01.

For the Pulsar search the settings were: maximum of two missed trypsin cleavages; PSM, Protein and Peptide FDR levels set to 0.01; scanning range set to 300-1800 m/z and Relative Intensity (Minimum) set to 5%; cysteine carbamidomethylation set as fixed modification and acetyl (N-term), deamidation (asparagine, glutamine), dioxidation (methionine, tryptophan), glutamine to pyro-Glu, oxidation of methionine and Phospho (STY) set as variable modifications. The databases used were *P. infestans* T30-4, downloaded from fungidb.org on 06-12-2023 (17,799 entries) modified with the sequences for mCherry and mCitrine, and *P. vulgaris* proteome, downloaded from uniprot.org (30,501 entries, UP000000226).

The resulting protein groups table was analysed in Perseus ^23^. From here on, the term “protein” is used interchangeably with “protein group” for simplicity. Proteins matching *Phaseolus vulgaris* proteins were removed (as potential contamination from ALB media), intensity values of the remaining protein groups for each sample normalised to the median intensity within the sample and subsequently Log_2_ transformed. Data from one of the four replicate experiments was excluded from downstream analysis at this point due to low protein coverage and identification compared to the remaining replicates.

Proteins classed as detected in the entire dataset were those present in at least two of all samples. Protein classed as detected within an individual fraction were those detected in at least two biological replicates of that fraction. The more stringent classification of present or absent within a fraction required a protein group to be present or absent in all three biological replicates of that fraction, respectively.

For comparison of protein abundance between fractions F2, F3, F4 and F6, proteins detected (as defined above) in all four fractions were used for the analysis. One-way ANOVA and Tukey’s HSD test (FDR <0.05 and q-value <0.05) were used to identify protein significantly differentially abundant between fractions. Proteins were then further filtered on -0.6 ≥ Log_2_(Fold Change) ≥ 0.6, equivalent to a Fold Change of at least 1.5 in the negative or positive direction, respectively, or for presence/absence. These filtered proteins were combined to form a set of high confidence proteins taken forward for further analysis.

For comparison of fraction F2 against F4, proteins detected (as defined above) in either F2 or F4 were used for the analysis. Two-sample t-test (FDR <0.05 and q-value <0.05) was used to identify proteins significantly differentially abundant between the fractions. Proteins were then further filtered on -0.6 ≥ Log_2_(Fold Change) ≥ 0.6, equivalent to a Fold Change of at least 1.5 in the negative or positive direction, respectively.

Euler’s diagrams, heatmaps, and volcano plots were drawn using the R packages ‘eulerr’ (version 7.0.2) ^24^, ‘gplots’ (version 3.2.0) ^25^, and ‘ggplot2’ ^26^, respectively.

### *In silico* prediction of protein cellular localisation and carbohydrate active enzymes

The predicted *P. infestans* T30-4 proteome was analysed using the DeepLoc 1.0 server (https://services.healthtech.dtu.dk/service.php?DeepLoc-1.0) ^27^ and the dbCAN3 server (https://bcb.unl.edu/dbCAN2/) ^28^ to predict protein cellular localisation and to identify potential carbohydrate active enzymes (CAZymes), respectively.

Four proteins were omitted from the DeepLoc anlaysis due to excessive length (PITG_01932, PITG_04003, PITG_04980 and PITG_12547). DeepLoc predictions for Plastids were combined with Mitochondrion, as *P. infestans* does not contain plastids. Counts for cellular localisation were done on the individual protein level (e.g., proteins within protein groups were counted individually) and statistical associations were assessed by Pearson’s Chi-squared test with Monte Carlo simulation (100,000 simulations) using the R package “chisq.posthoc.test” with Bonferroni-Holm correction for multiple testing ^29^.

### Identification of signal peptide containing proteins and RXLRs

To identify the presence of signal peptide containing proteins, protein lists were searched against a reference list of 2,365 predicted signal peptide-containing proteins within the *P. infestans* genome. This reference list was generated by combining predicted signal peptide containing protein lists from the FungiDB website (https://fungidb.org/fungidb/app/, release 68), Meijer *et al*. ^30^, and Raffaele *et al.* ^31^ (Supplementary Dataset 2A). The presence of RxLR effector proteins was determined by sub-setting proteins with the term ‘RxLR’ within the ‘ProteinDescription’ field.

### Gene Ontology analysis

Gene Ontology (GO) analysis was performed on the individual protein level in ShinyGO v0.81 ^32^ available at https://bioinformatics.sdstate.edu/go using all detected proteins as the background set, with species database set to *Phytophthora infestans* genes ASM14294v1. The analysis was set to return the top 10 pathways for Biological Process, with a minimum pathway size of 10. All other settings were left at default, with FDR set to 0.05. For lollipop diagrams of enriched GO terms, top GO terms with a minimum of five proteins identified were displayed. In some cases where redundancy between GO terms was high, proteins in the top three GO terms were collapsed into a single non-redundant list. Interpro and Pfam protein domain descriptions obtained from FungiDB (https://fungidb.org/fungidb/app/, release 68) were used to infer protein descriptions/function where annotation was lacking.

## Results

### Generation of a dual expression vector enables visualisation and tracking of fluorescently-tagged secreted proteins

Given the lack of verified vesicle markers in *P. infestans,* we instead used two effector proteins as markers for secretory proteins/vesicle cargo: PITG_04314, a representative of the RXLR effectors, and the pectin esterase PITG_01029, a host cell wall modifying enzyme within the apoplastic effector category ^11^.

To track the cellular location of these two effector proteins, a dual expression vector (pDual_mCherry_mCitrine, Fig. S1) was constructed and used to simultaneously express PITG_04314-mCherry with PITG_01029-mCitrine in *P. infestans*. Mycelia of the dual transformant was grown *in vitro* in liquid culture and western blot analysis of protein extracted from the mycelia and conditioned medium confirmed both tagged proteins were expressed and secreted from mycelia (Fig. 1A). PITG_01029-mCitrine appeared to run to an approximate size of 100 kDa, larger than the expected 61.5 – 63 kDa. To determine if this was due to glycosylation, a deglycosylation assay was performed which confirmed this size shift was due to posttranslational modification (Fig. S2). PITG_04314-mCherry ran as a doublet in the expected size range of 42 – 44.5 kDa, which remained unaltered after treatment with the deglycosylation enzyme mix. This suggests presence of the doublet was not due to glycosylation state and may instead be due to proteolytic processing of the protein ^33^ (Fig S2).

**Figure 1.**
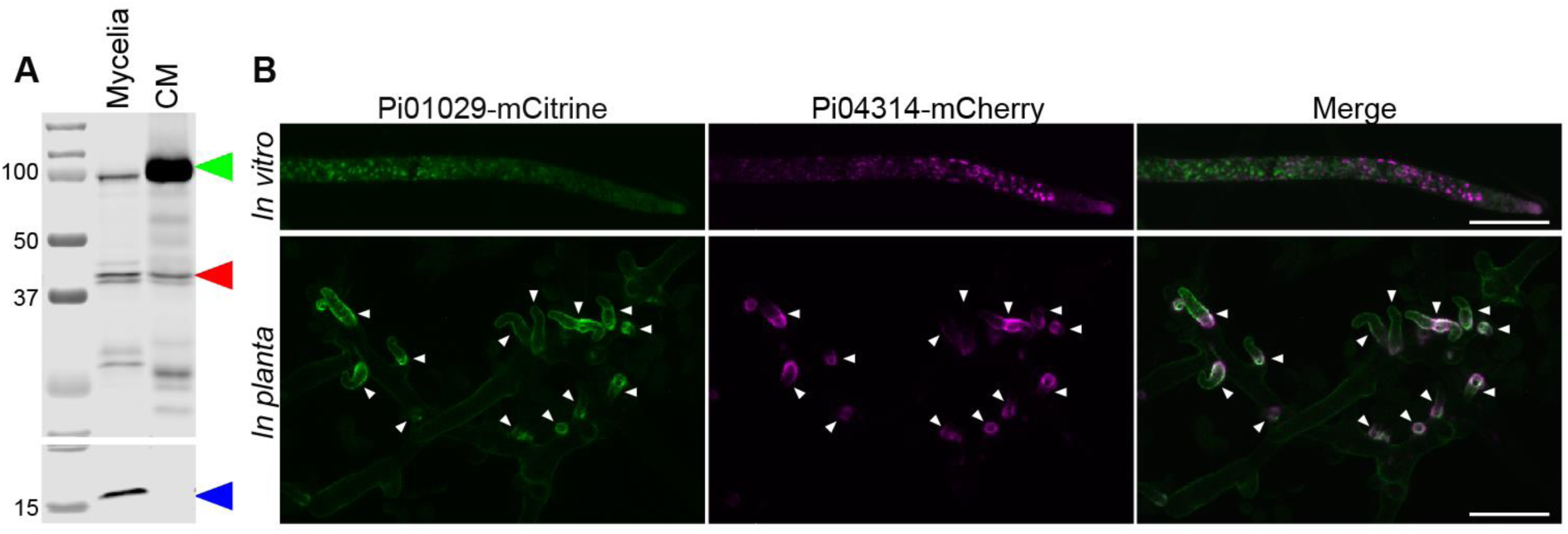
Simultaneous localisation of PITG_04314-mCherry and PITG_01029-mCitrine expressed from a dual expression vector in *P. infestans*. (A) Western blot analysis of *in vitro* grown mycelia and the corresponding conditioned media (CM). Green arrowhead: PITG_01029-mCitrine, red arrowhead: PITG_04314-mCherry, blue arrowhead: histone. Sizes of marker proteins (kDa) are indicated. (B) Confocal microscopy of mycelia grown *in vitro* (upper panel) and *in planta* (lower panel). White arrowheads indicate location of haustoria. Scale bars indicate 20 µm.

Confocal microscopy of the mycelia grown *in vitro* confirmed the correct expression and folding of the fluorescent tags, with both tagged proteins appearing to localise to punctate structures (Fig. 1B, upper panel). Confocal imaging of the transformant *in planta* during infection of *Nicotiana benthamiana* leaves shows both PITG_04314-mCherry and PITG_01029-mCitrine accumulating at the haustoria (Fig. 1B, lower panel).

### Buoyant density ultracentrifugation facilitates robust and reproducible isolation of *P. infestans* intracellular vesicles

As both PITG_04314-mCherry and PITG_01029-mCitrine were confirmed to be secreted from mycelia, we set out to isolate the intracellular vesicles in which these proteins are trafficked along the secretory pathway. Using the dual transformant, we isolated intracellular vesicles from *in vitro* mycelia, using cushioned bottom-loaded density gradient ultracentrifugation (Fig. 2). With this technique, vesicles are concentrated onto a high density iodixanol cushion without pellet formation, which increases recovery and preserves the integrity and morphology of the vesicles ^34^. The concentrated vesicles were then bottom-loaded and overlaid with gradient medium (iodixanol) of progressively lower density. During ultracentrifugation, the vesicles migrate up the gradient to a point at which their density matches that of the surrounding gradient medium, separating the vesicles based on buoyancy.

**Figure 2.**
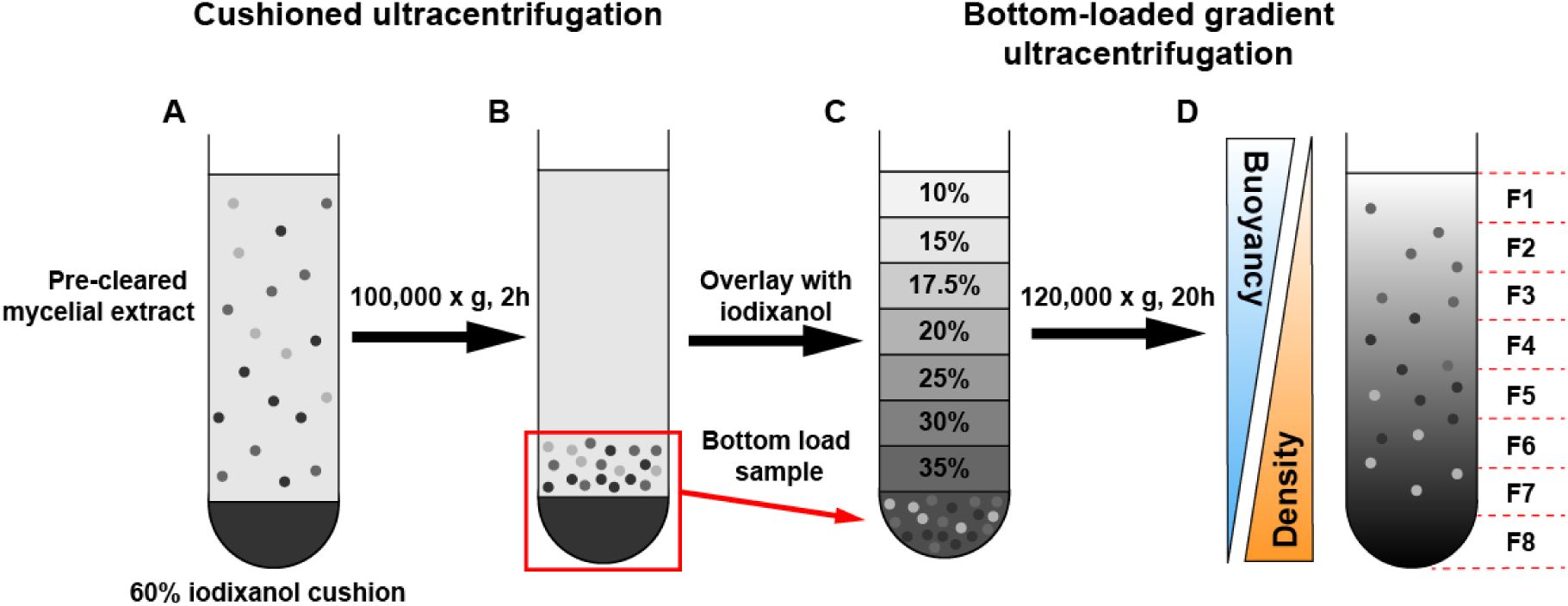
Overview of cushioned-bottom loaded iodixanol gradient ultracentrifugation method used to isolate *P. infestans* intracellular vesicles based on buoyancy. (A) Pre-cleared mycelial extract containing intracellular vesicles in isolation buffer were loaded on top of a 60% iodixanol cushion and centrifuged at 100,000 × g for 2 h. (B) After centrifugation, the vesicles concentrate above the cushion. (C) The iodixanol cushion and 1 ml above the cushion containing the concentrated vesicles were mixed, bottom loaded into a new tube, then overlayed with layers of iodixanol with progressively lower densities to form a discontinuous gradient. (D) After centrifugation at 120,000 × g for 20 h, the gradient becomes continuous, with vesicles migrating up the gradient based on buoyancy, with denser material remaining near the bottom of the tube. Eight fractions were taken starting from the top of the tube to the bottom for analysis (fractions F1-F8).

Replicate experiments revealed the gradients to be reproducible in terms of floatation of PITG_04314-mCherry and PITG_01029-mCitrine up the gradient and density measurements of the fractions. PITG_04314-mCherry and PITG_01029-mCitrine consistently migrated up to regions of lower density, predominantly migrating to fractions F2, F3 and F4 (Fig. 3A, C), corresponding to average densities of 1.111 ± 0.013 g/mL, 1.128 ± 0.009 g/mL and 1.150 ± 0.012 g/mL, respectively (Fig. 3B). Total protein staining showed migration of proteins up the gradient, with a clear distinction in protein profiles between the lower density fractions F1-F4 and the higher density fractions F5-F8 (Fig. S3). Transmission electron microscopy (TEM) imaging of fractions F2, F3, and F4 showed the presence of vesicular structures, whereas the higher density fraction F6, containing little PITG_04314-mCherry and PITG_01029-mCitrine, showed a near absence of vesicle-like structures (Fig. 3D).

**Figure 3.**
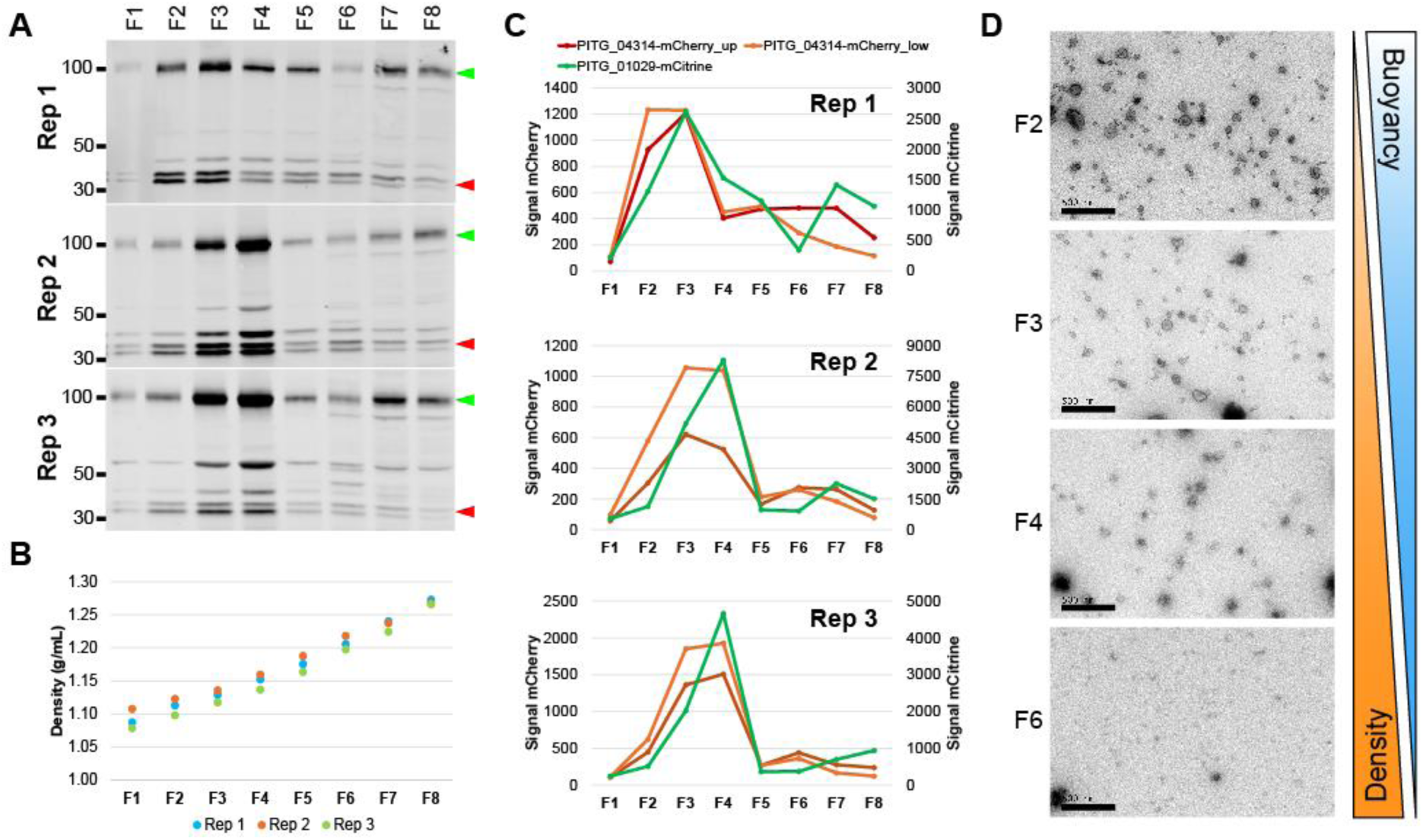
PITG_04314-mCherry and PITG_01029-mCitrine migrate to fractions of lower density within the gradient, coinciding with the presence of vesicle-like structures. (A) Western blot of fractions taken from three independent gradients. Green arrowhead: PITG_01029-mCitrine, red arrowhead: PITG_04315-mCherry. Sizes of marker proteins (kDa) are indicated. (B) Density measurements calculated from OD 340 nm readings of the fractions shown in (A). (C) Quantification of band intensities for PITG_04314-mCherry and PITG_01029-mCitrine shown in (A). PITG_04314-mCherry appears as a doublet; intensity of the upper (‘PITG_04314-mCherry_up’) and lower (‘PITG_04314-mCherry_low’) were quantified separately. (D) Transmission electron microscopy (TEM) images of selected fractions. Scale bars: 500 nm.

### Disruption of vesicle integrity prevents differential protein migration within the gradient

Addition of detergent such as Triton X-100 to the sample prior to bottom-loading of the gradient is expected to disrupt any membrane-bound structures such as vesicles, causing release of any protein cargo and disrupting floatation.

Without Triton X-100 pretreatment, PITG_04314-mCherry and PITG_01029-mCitrine migrated predominantly to fractions F2, F3 and F4 as expected (Fig. 4A, C). On the other hand, sample pretreated with Triton X-100 run in parallel showed a measurable shift in the migration of PITG_04314-mCherry and PITG_01029-mCitrine, with accumulation towards the bottom/denser portions of the gradient (Fig. 4B, D). Pretreatment with Triton X-100 affected protein migration within the gradient in general, as total protein staining of western blots showed increased accumulation of total protein in fractions F5-F8 with the absence of detectable protein staining in fractions F1-F4 (Fig. S4). This suggests that Triton X-100 disrupts intracellular vesicles in general. TEM of fractions F2 and F3 in the untreated sample revealed the presence of vesicular structures, which were absent in the sample pretreated with Triton X-100 (Fig. 4E, F).

**Figure 4.**
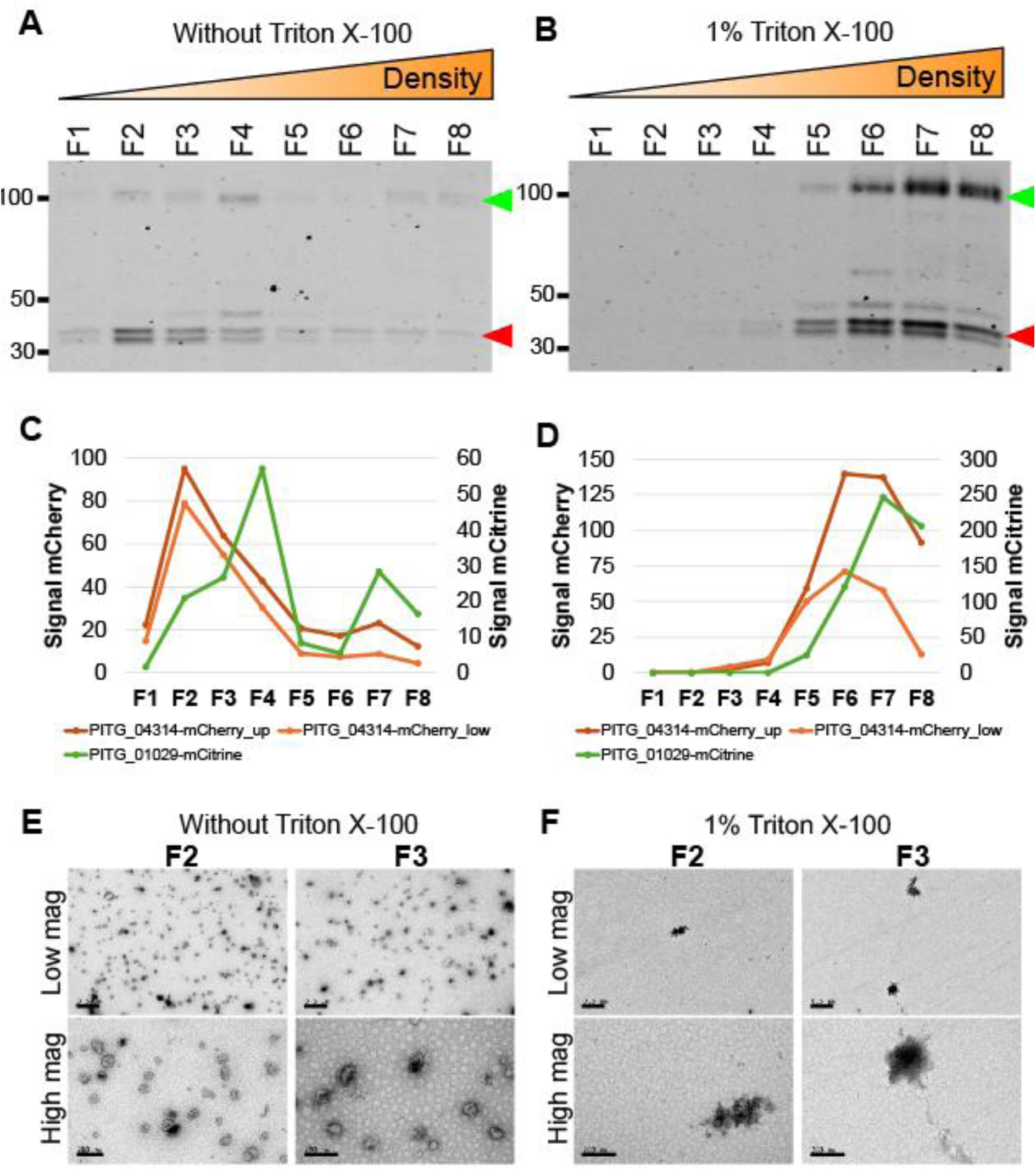
Preincubation of samples with 1% Triton X-100 prior to bottom-loading of the gradient prevents migration of PITG_04314-mCherry and PITG_01029-mCitrine up the gradient, coinciding with the absence of vesicle-like structures. (A) Western blot showing position of PITG_04314-mCherry and PITG_01029-mCitrine within the fractions of a gradient in the absence of Triton X-100. Sizes of marker proteins (kDa) are indicated. Green arrows: PITG_01029-mCitrine, red arrows: PITG_04314-mCherry. (B) Western blot showing position of PITG_04314-mCherry and PITG_01029-mCitrine within a gradient on pre-incubation of the sample with 1% Triton X-100 prior to bottom-loading. (C) Quantification of band intensities for PITG_04314-mCherry and PITG_01029-mCitrine shown in (A) (without Triton X-100). (D) Quantification of band intensities for PITG_04314-mCherry and PITG_01029-mCitrine shown in (B) (with 1% Triton X-100). (E) Transmission electron microscopy (TEM) images of fractions F2 and F3 in the absence of Triton X-100 and (F) in the presence of 1% Triton X-100. High magnification scale bars: 200 nm, Low magnification scale bars: 0.5 µm.

These results suggest that PITG-04314-mCherry and PITG-01029-mCitrine are likely trafficked within the cell in vesicles and that our protocol enables isolation of these vesicles whilst preserving their integrity.

### Presence of vesicular structures in buoyant fractions correlates with differences in protein abundance profiles

As PITG_04314-mCherry and PITG_01029-mCitrine migrated predominantly to the low-density fractions F2, F3 and F4, these buoyant fractions were analysed to identify proteins associated with *P. infestans* intracellular vesicles. Since the high-density fraction F6 was found to be devoid of vesicle-like structures, we included this fraction to account for potential contaminants/proteins not associated with intact vesicles.

Across all fractions, a total of 6,685 *P. infestans* protein groups were detected (i.e., present in at least two samples) (Supplementary Dataset 1A). Hierarchical clustering based on protein abundance profiles across fractions revealed the buoyant fractions F2, F3 and F4 to be more similar to each other than to the denser fraction F6 (Figure 5A).

**Figure 5.**
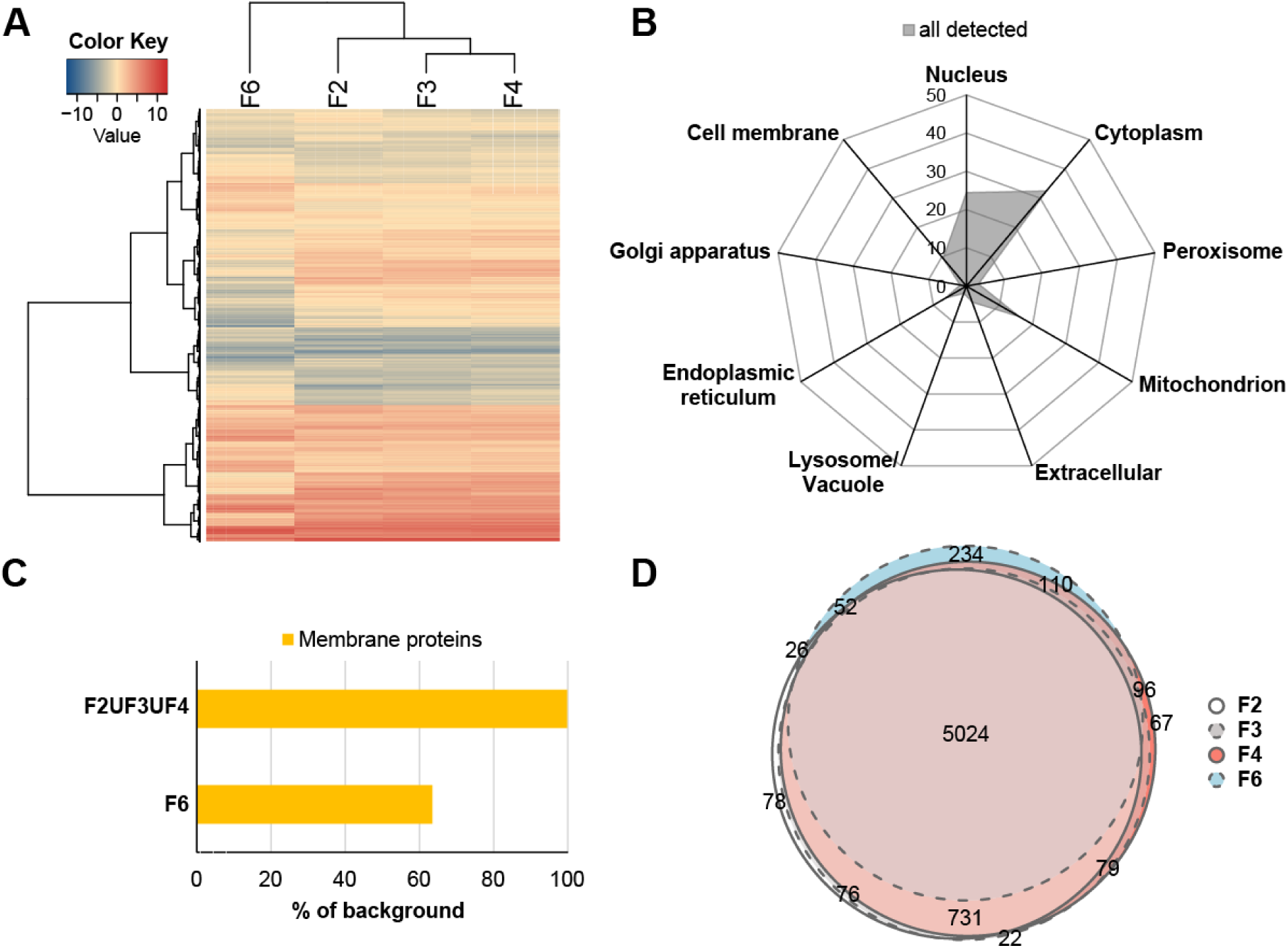
Overview of Data-Independent Acquisition mass spectrometry data of fractions F2, F3, F4 and F6 from three biological replicate experiments. (A) Heatmap displaying hierarchical clustering of fractions based on similarities in protein intensity for the 6,685 protein groups detected by mass spectrometry run in Data-Independent Acquisition (DIA) mode for each fraction. (B) Radar chart showing DeepLoc predicted cellular association of the detected proteins as a percentage of total proteins detected. (C) Number of membrane proteins detected in at least one buoyant fraction (F2ՍF3ՍF4) and F6 as a percentage of the 1,626 predicted membrane proteins detected in the entire dataset. (D) Euler diagram showing overlap in proteins detected (present in at least two reps) for fractions F2, F3, F4 and F6. The overlapping 5,024 proteins present in all four fractions were taken forward for comparison of protein abundance between fractions.

Prediction of cellular localisation of the detected proteins revealed the most abundant proteins to be associated with the cytoplasm (32.5%), nucleus (24.4%) and mitochondrion (15.9%). (Figure 5B). Lower abundance proteins were associated with cell membrane (9.5%), endoplasmic reticulum (6.3%), extracellular (4.5%), peroxisome (3.2%), lysosome/vacuole (2.0%) and Golgi apparatus (1.7%). Classification of proteins as either membrane or soluble revealed all but three (PITG_13657, PITG_06609 and PITG_15810) of the 1,626 detected membrane proteins could be found in at least one of the buoyant fractions (Figure 5C). In contrast, only 594 of the total detected membrane proteins (36.5%) were detected in F6 (Figure 5C).

Within individual fractions, the number of proteins present (i.e., identified in at least two biological replicates of the fraction) were 6,057, 6,078, 6,199 and 5,592 proteins for fractions F2, F3, F4 and F6 (respectively), with an overlap of 5,024, suggesting high similarity regarding the identity of proteins present within each fraction (Figure 5D). These observations suggest differences in protein abundance, rather than presence or absence, as the main factor determining differences between buoyant and dense fractions. Indeed, analysis of protein presence/absence revealed 283 proteins that are exclusively associated with all three buoyant fractions (Supplementary Figure S5a), and only three proteins exclusively found in F6 (Supplementary Figure S5b).

Given that we were interested in proteins enriched in the buoyant fractions associated with intracellular vesicles, enrichment analysis focussed on comparison of fractions F2, F3 and F4 versus the denser fraction F6, which was shown to be devoid of vesicles. The flowchart in Figure 6 summarises the workflow used for protein enrichment analysis.

**Figure 6.**
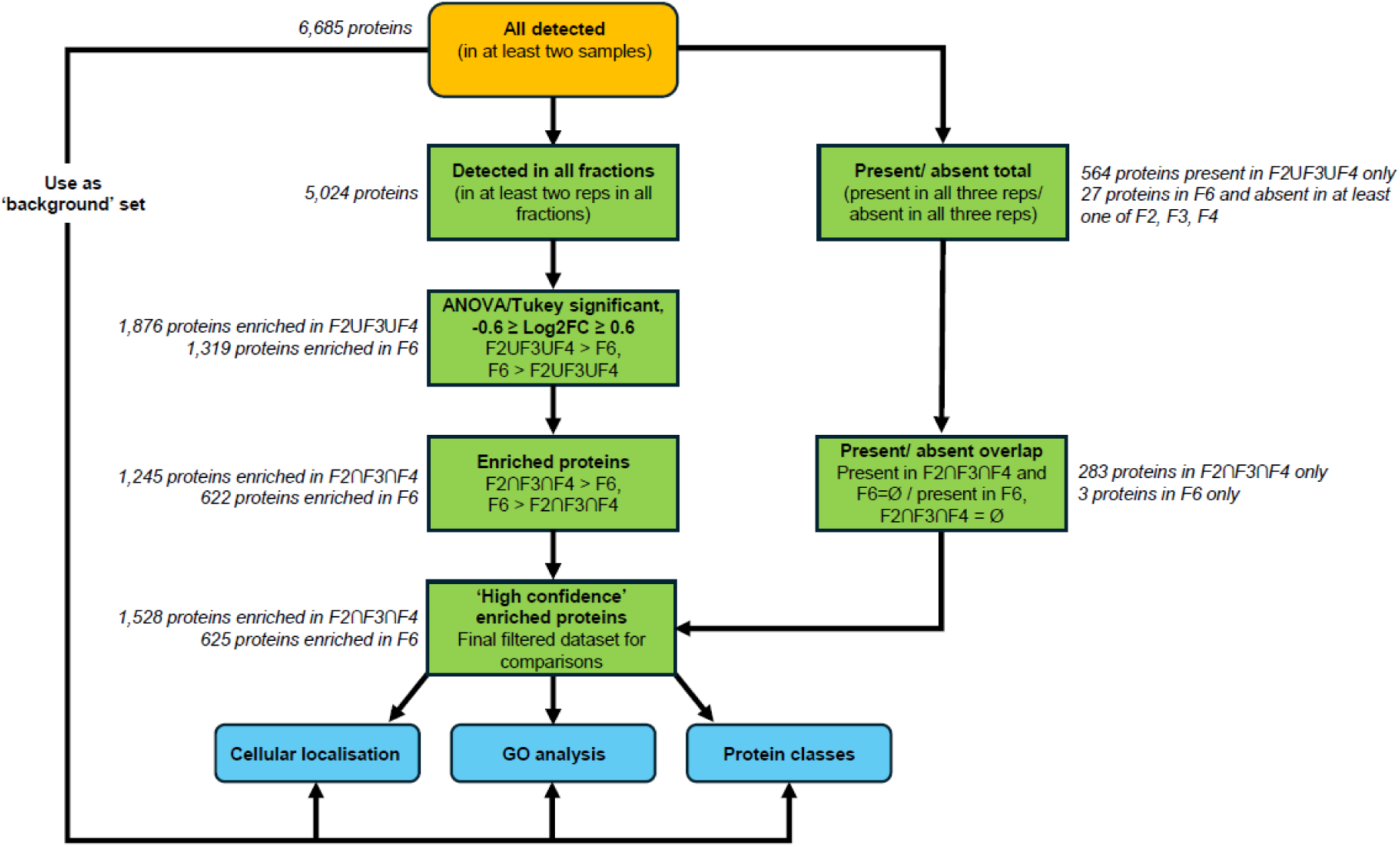
Schematic of data analysis workflow. A set of high confidence proteins for enrichment analysis were derived from filtering for proteins based on presence/absence and Log_2_ Fold-change (Log2FC). ‘F2ՍF3ՍF4’: set of proteins representing the union of proteins detected in F2, F3 and F4 (i.e., present in at least two reps for at least one of the fractions), ‘F2∩F3∩F4’: set of proteins representing the intersect of proteins detected in F2, F3 and F4 (i.e., present in at least two reps for all fractions), ‘F6 = Ø’: where F6 is the empty set in relation to presence/absence, ‘F2∩F3∩F4 = Ø’: where F2∩F3∩F4 is the empty set in relation to presence/absence.

### Buoyant fractions F2, F3 and F4 are enriched for a common set of proteins

The 5,024 proteins common between all fractions were further analysed for differential abundance between fractions. One-way ANOVA (FDR 0.05, q-value <0.05 with Tukey’s HSD test (FDR 0.05)) was performed to identify proteins which differed significantly between the buoyant fractions and fraction F6, returning a total of 3,320 proteins. Log_2_Fold-Change (Log_2_FC) was calculated for each protein as the ratio of median normalised intensities for each buoyant fraction against fraction F6. The proteins were then further filtered based on a Log_2_FC cutoff of -0.6 ≥ Log_2_FC ≥ 0.6 to obtain the differentially abundant proteins within the fractions; those more abundant in at least one of the buoyant fractions F2, F3 and F4 (F2ՍF3ՍF4 > F6, Log_2_FC ≥ 0.6), of which there were 1,876, and those more abundant in the high density fraction F6 compared to at least one buoyant fraction (F6 > F2ՍF3ՍF4, Log_2_FC ≤ -0.6), of which there were 1,319.

Within the 1,876 proteins enriched in at least one of the buoyant fractions, 1,245 intersected or were more abundant in all three fractions over F6 (F2ՈF3ՈF4 > F6, Log_2_FC ≥ 0.6) (Supplementary Figure S5C). These 1,245 proteins were combined with the 283 proteins that were present in all three buoyant fractions and absent in F6 to give a set of 1,528 high confidence proteins potentially associated with intracellular vesicles.

Of the 1,319 proteins identified as more abundant in F6 over at least one buoyant fraction, 622 of these were enriched in F6 over all three buoyant fractions (F6 > F2ՈF3ՈF4, Log_2_FC ≤ -0.6) (Supplementary Figure S5D). These 622 proteins were combined with the three proteins present exclusively in F6 to give a set of 625 high confidence proteins enriched in fraction F6, representing potential contaminant proteins not associated with intracellular vesicles.

With the 1,528 (Supplementary Dataset 1B) and 625 (Supplementary Dataset 1C) high confidence proteins enriched in the buoyant fractions and F6, respectively, we proceeded to classify the types of proteins within these two datasets, in the context of the full background set of all 6,685 detected proteins.

### Buoyant fractions are enriched for membrane proteins and proteins that enter the conventional secretory pathway

As vesicles are membrane bound structures, fractions enriched for these structures would be expected to be enriched for membrane proteins as opposed to soluble proteins. To determine if this were the case, the proteins present in the fractions were classed as either membrane or soluble through DeepLoc prediction ^27^. Soluble proteins were further subdivided into those with a signal peptide (+SP), which are predicted to enter the conventional ER-to-Golgi secretory pathway, and those without (-SP) (Figure 7A). In the background set of all detected proteins, almost a quarter were predicted to be membrane localised (24.3%, 1,626 proteins), with the remaining three quarters consisting of soluble proteins. Compared to this, the buoyant fractions were enriched in membrane proteins (62.2%, 952 proteins), whilst F6 was clearly depleted (1.3%, 8 proteins). Although F6 mainly consisted of soluble proteins, making up a total of 98.7% of all proteins present in this fraction compared to 37.7% in the buoyant fractions, only 1.9% (12 proteins) were both soluble and contained a predicted SP, compared to 10.4% (159) of soluble proteins with a signal peptide in the buoyant fractions (Supplementary Dataset 2A). The background set of all detected proteins consisted of 6.1% (409) soluble proteins with signal peptide.

**Figure 7.**
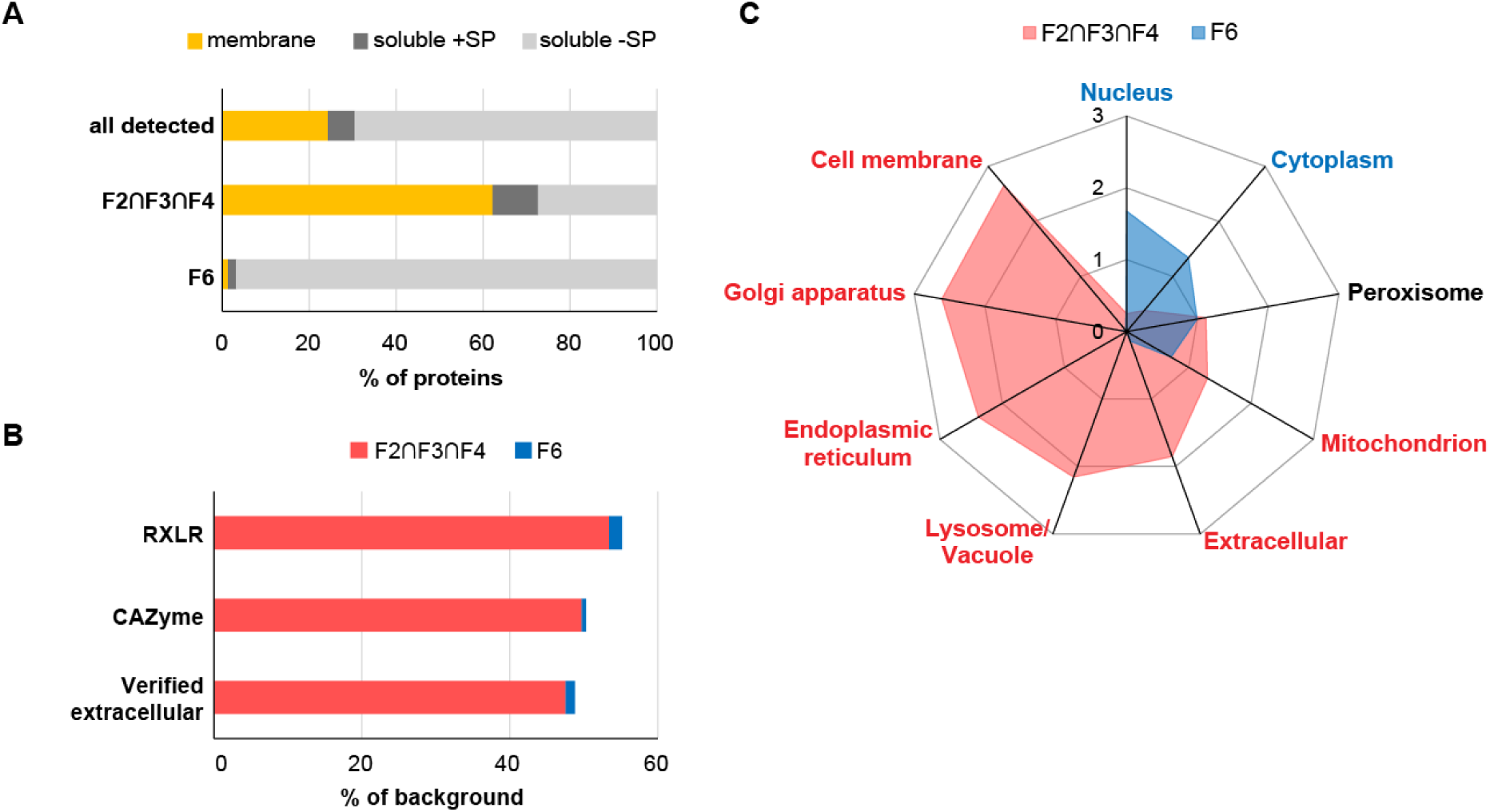
Classification and associated cellular localisation of proteins enriched in fractions F2, F3, F4 and F6. (A) Percentage of proteins predicted membrane associated or soluble found in all three buoyant fractions (F2∩F3∩F4), and within F6. The predicted soluble proteins were further subdivided into those with (‘soluble +SP’) and without (‘soluble -SP) a signal peptide. The proportion of each class of protein for the background set of all 6,685 detected proteins (‘all detected’) is also shown for comparison. (B) Number of RXLR and CAZymes associated with the buoyant fractions and with F6 shown as a percentage of the total number of these proteins in the background set of all detected proteins. The number of ‘verified extracellular’ proteins were determined by comparison to a list of proteins previously identified as secreted into the extracellular space by *P. infestans.* See main text for details. (C) Radar chart showing percentage of proteins associated with each cellular compartment (predicted by DeepLoc) as a ratio of the percentage found in the background set (fold-enrichment) for the buoyant fractions (F2∩F3∩F4, pink) and F6 (blue). Cellular localisation labelled in red showed significant association with the buoyant fractions, and those labelled in blue showed significant association with F6. Labels in black show no significant association. Significance was determined by Monte Carlo Chi-squared test (p<0.001).

### Effector proteins are associated with buoyant fractions

Since the RXLR effector protein PITG_04314 and the apoplastic effector/pectin esterase PITG_01029 were shown to be associated with the vesicle-rich buoyant fractions, we set out to determine if this was a general trend in our dataset. Of the annotated 596 RXLR effectors in the *P. infestans* database, 58 were detected in our dataset, 31 (53.4%) of which were enriched within the buoyant fractions, and only one (1.7%) enriched within F6 (Figure 7B). For carbohydrate active enzymes (CAZymes), which include pectinesterases, 505 were predicted to be encoded in the *P. infestans* genome; 181 of these were detected in our dataset, of which 90 (49.7%) were enriched in the buoyant fractions, and one (0.5%) enriched within F6 (Figure 7B).

These findings together suggest buoyant fractions enrich for membrane proteins and for signal peptide containing soluble proteins which enter the secretory pathway, such as RXLR effectors and CAZymes, whereas F6 is enriched for non-secretory soluble proteins.

To determine whether our data set captured *bona fide* extracellular secreted proteins, we compared our data set to a published *P. infestans* extracellular secretome. Meijer *et al*. ^30^ previously profiled the extracellular proteome of *P. infestans* mycelia grown in seven different types of media and collectively identified 283 extracellular *P. Infestans* proteins. The majority of the secreted proteins found were those involved in defence against oxidative stress, pathogen associated proteins such as effectors and elicitins, and various proteases (aspartic, cysteine, and metalloproteases). Of these 283 proteins, over half (164 proteins) were detected in our proteomics data set, of which 47.5% (78 proteins) were enriched within the buoyant fractions, with only 1.2% (two proteins) enriched within F6 (Figure 7B, Supplementary Dataset 2C).

### Buoyant gradient ultracentrifugation separates proteins associated with different cellular compartments

To obtain an overview of the intracellular sources of the proteins found within the fractions, the enriched proteins were sub-divided into their predicted cellular localisations.

To refine the classes of proteins found in the fractions, cellular localisation of the proteins as predicted by DeepLoc was compared between the buoyant fractions and F6. This comparison revealed significant association between the type of fraction (buoyant vs dense) and the predicted cellular localisations of the proteins present, except for proteins associated with peroxisomes which was similar between both types of fractions (Figure 7C). Proteins associated with the nucleus (43.5%) and cytoplasm (40.9%) represented the most abundant types of proteins in F6. Proteins associated with cell membrane, Golgi apparatus, and the extracellular compartment collectively formed less than 1% of the enriched proteins in F6, and no proteins associated with lysosome/vacuole or endoplasmic reticulum were identified as enriched in F6. In complete contrast, the buoyant fractions were depleted in cytoplasmic (12.4%) and nuclear proteins (6.1%) but were enriched for those associated with cell membrane (25.1%), endoplasmic reticulum (15%), extracellular (8.3%), Golgi apparatus (4.4%) and lysosome/vacuole (4.3%), totalling 57.2% of the total enriched proteins.

The buoyant fractions were also more enriched in mitochondrial proteins compared to F6 (20.7% vs 11.4%, respectively). However, the nature of the mitochondrial proteins enriched differed – 66.1% of mitochondrial proteins enriched in the buoyant fraction were predicted to be membrane-associated, whereas only 7% of mitochondrial proteins enriched in F6 had this property, with the majority predicted to be soluble (Supplementary Figure S6).

### Gene Ontology analysis reveals enrichment for proteins associated with membranes and vesicle trafficking in buoyant fractions

Since the data suggested the buoyant fractions and the denser F6 fraction differ in the general types of proteins enriched and their subcellular origins, we used Gene Ontology (GO) analysis to determine whether these two sets of proteins also differed in their associated biological functions.

Both the 1,528 proteins enriched in the buoyant fractions and the 625 proteins enriched in F6 were compared against the background set of all 6,685 detected proteins to look for enrichment of GO terms belonging to the Biological Process category.

‘Transmembrane transport’ (GO:0055085, 203 proteins), ‘Establishment of localisation’ (GO:0051234, 322 proteins), and ‘Localisation’ (GO:0051179, 327 proteins) were identified as the top three GO terms for the buoyant fractions (Figure 8A, Supplementary Dataset 3A). For F6, ‘Gene expression’ (GO:0010467, 221 proteins), ‘Cellular nitrogen compound metabolic process’ (GO:0034641, 265 proteins), and ‘Translation’ (GO:0006412, 268 proteins) were identified as the top three GO terms (Figure 8B, Supplementary Dataset 4A).

**Figure 8.**
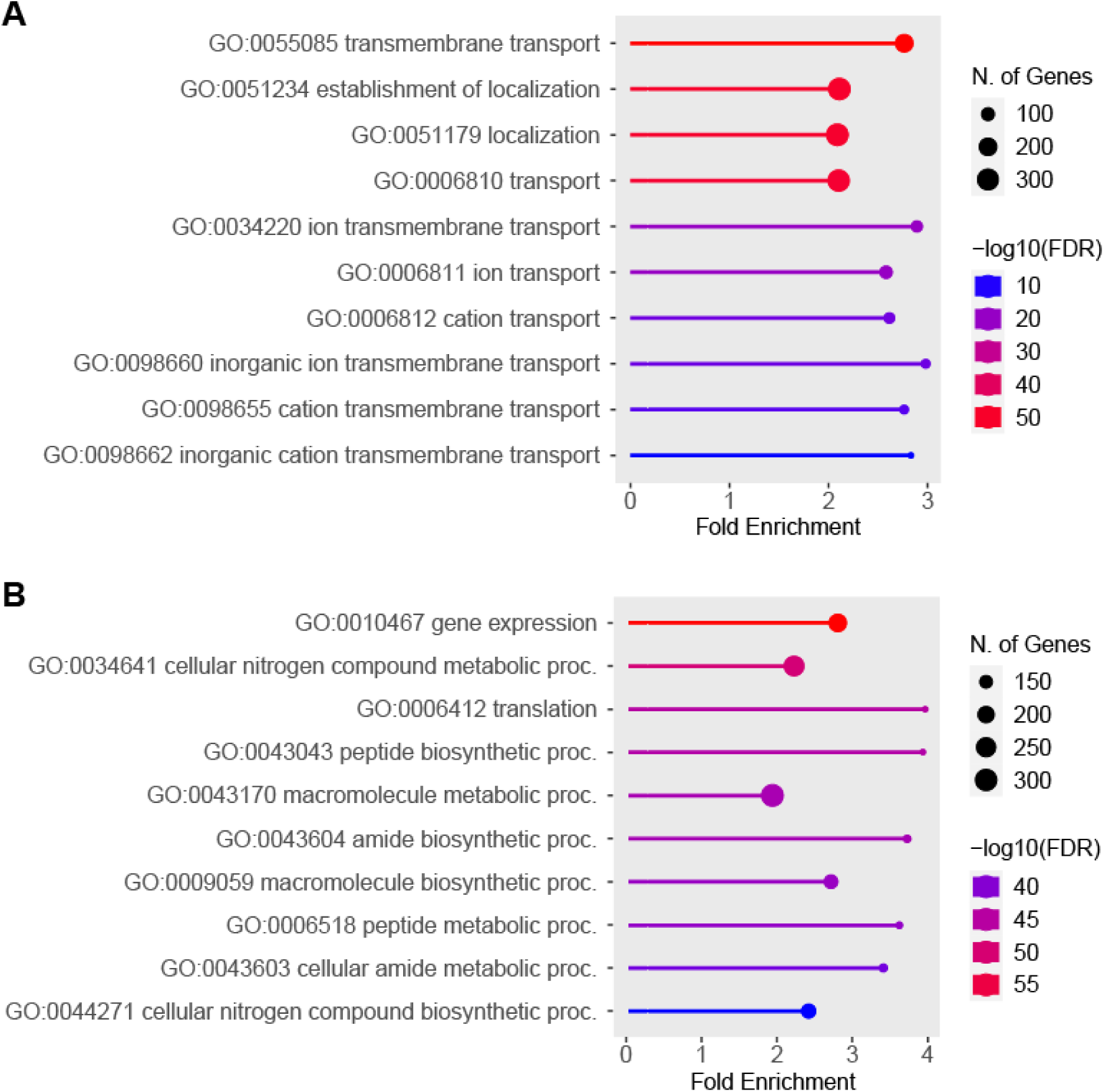
Top ten Gene Ontology terms enriched in the buoyant fractions and F6 as assessed by ShinyGO. (A) Biological Process GO terms enriched in the buoyant fractions F2, F3 and F4. (B) Biological Process GO terms enriched in fraction F6.

Since proteins can be assigned more than one GO term, with large redundancy between some terms, the proteins contained within the top three GO terms were collapsed to give a single non-redundant list for further analysis. For the buoyant fractions, this resulted in a collapsed set of 327 proteins encompassing proteins from all three GO terms (Supplementary Dataset 3B), and for F6, the collapsed set resulted in a list of 268 proteins (Supplementary Dataset 4B).

Interrogation of the proteins included in the top three GO terms enriched in the buoyant fraction included a range of membrane trafficking proteins, such as 16 Ras-associated binding (Rab) proteins, eight vacuolar protein sorting associated (Vps) proteins, 17 soluble *N*-ethylmaleimide-sensitive factor attachment protein receptor (SNARE) proteins and proteins related to SNARE function, six Golgi Dynamics (GOLD) domain-containing proteins, and various other proteins involved in vesicle trafficking (Supplementary Dataset 3B).

Membrane transporters/proton pumps were also enriched in the buoyant fractions, with proteins forming V-ATPases (six proteins), ABC transporters (41 proteins), MSF transporters (33 proteins), and P-type ATPases (18 proteins) identified. In addition to their role in protein transport across membranes, V-ATPases have also been implicated in membrane fusion events, independent of their ATPase activity ^35^. Figure 9 summarises the cellular localisations of the enriched proteins.

**Figure 9.**
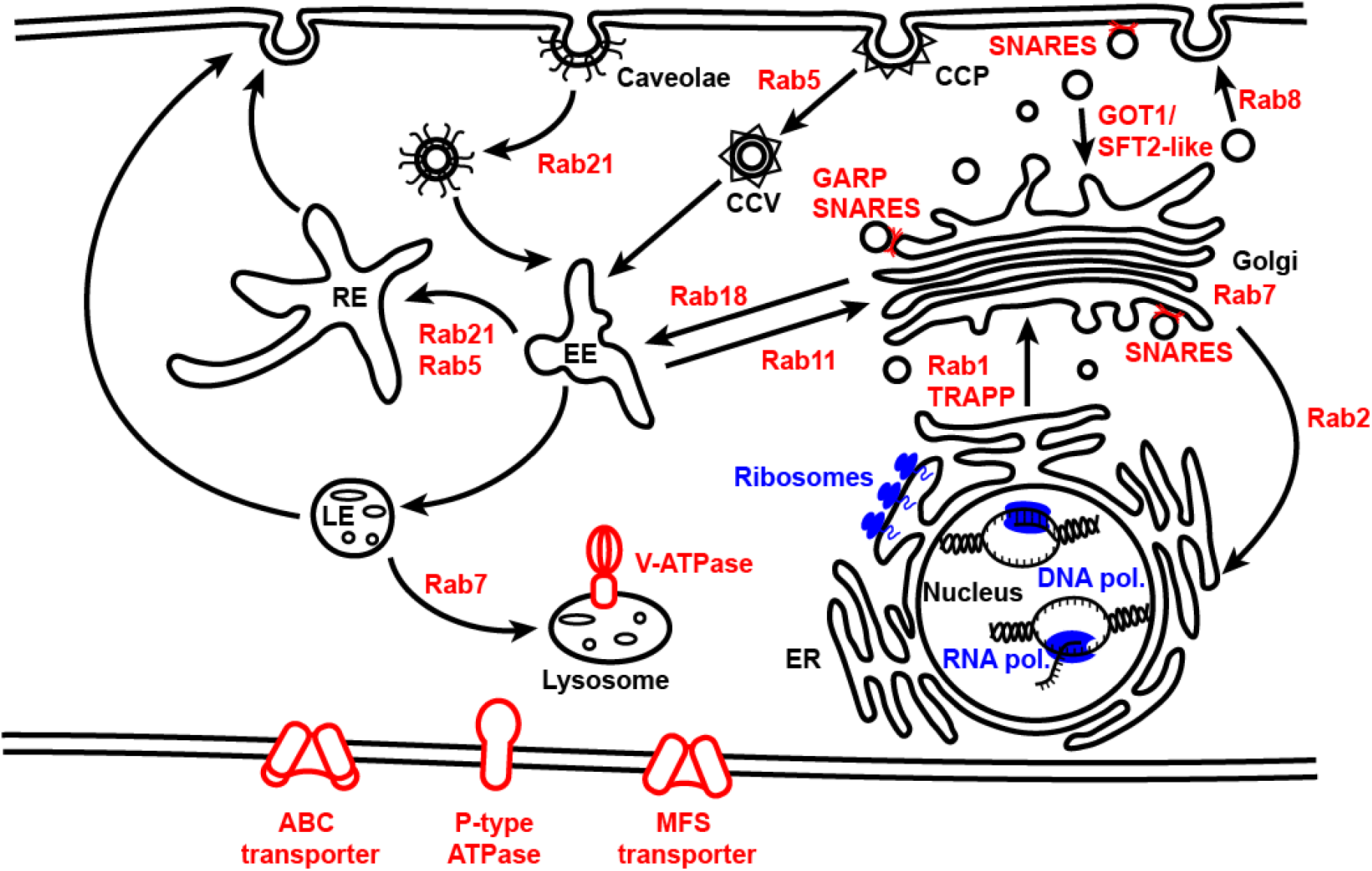
Overview of cellular localisation of enriched proteins from GO analysis. Proteins enriched in buoyant fractions (red) and in F6 (blue) are associated with distinct cellular compartments. CCP: clathrin-coated pit, CCV: clathrin-coated vesicle, RE: recycling endosome, EE: early endosome, LE: late endosome, ER: endoplasmic reticulum.

In contrast, the proteins enriched in the dense F6 fraction were dominated by ribosomal subunits and ribosome-associated proteins, DNA and RNA polymerases/proteins involved in transcription, translation, DNA repair and replication, and ribonuclear proteins (Supplementary Dataset 4B).

### Buoyant density ultracentrifugation separates intracellular vesicles with differing protein content

To determine whether the present protocol was able to separate proteins based on their association with intracellular vesicles of differing buoyancy characteristics, we compared protein content of the buoyant fractions F2 (1.111 ± 0.013 g/mL) and F4 (1.150 ± 0.012 g/mL). We chose these two fractions for comparison as they represent the extreme ends of the buoyant fractions analysed here in terms of density.

Analysis of differential protein abundance identified 1,330 and 1,909 proteins which were significantly more abundant in F2 over F4 (F2>F4) and F4 over F2 (F4>F2), respectively (Figure 10A). Amongst the differentially abundant proteins was the tetraspanin-like PiTET3 (PITG_12224), and the MARVEL protein PiMDP2 (PITG_13661), which were significantly more abundant in F2. In addition, 44 and 75 proteins were found to be exclusive to F2 and F4, respectively. This gave a combined total of 1,374 and 1,984 proteins more associated with F2 or F4, respectively (Supplementary Dataset 5).

**Figure 10.**
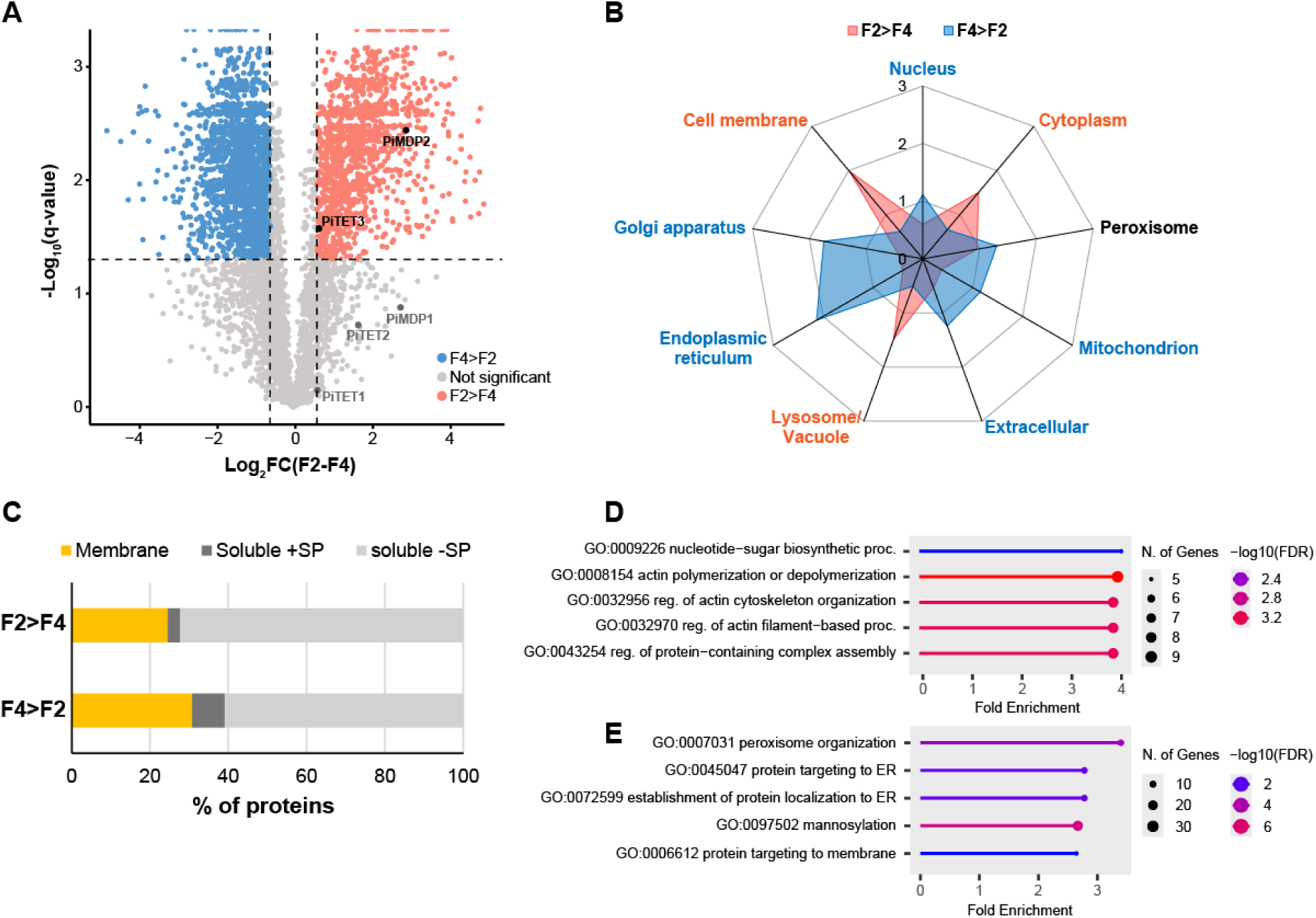
Comparison of proteins present in buoyant fractions F2 and F4. (A) Volcano plot showing proteins differentially abundant between F2 and F4, defined as those with -0.6 ≤ Log2FC(F2-F4) ≥ 0.6 and q-value <0.05. The positions of three *P. infestans* tetraspanins (PiTET1 (PITG_03954), PiTET2 (PITG_03950) and PiTET3 (PITG_12224)), and two MARVEL proteins (PiMDP1 (PITG_13660) and PiMDP2 (PITG_13661)) are highlighted in the plot. (B) Radar chart showing percentage of proteins associated with each cellular compartment (predicted by DeepLoc) as a ratio of the percentage found in the background set (fold-enrichment) for proteins enriched in F2 (F2>F4, red) and proteins enriched in F4 (F4>F2, blue). Cellular localisations labelled in pink showed significant association with fraction F2, and those labelled in blue showed significant association with F4. Labels in black show no significant association. Significance was determined by Monte Carlo Chi-squared test (p<0.001). (C) Percentage of proteins predicted membrane associated or soluble found enriched in F2 and F4. The predicted soluble proteins were further subdivided into those with (‘soluble +SP’) and without (‘soluble -SP) a signal peptide. (D) Top five GO terms for Biological Process enriched in fraction F2 over F4. (E) Top five GO terms for Biological Process enriched in fraction F4 over F2.

DeepLoc prediction of cellular localisations indicated proteins enriched in F2 had a higher association with the cell membrane, lysosomes and cytoplasm, whereas proteins more abundant in F4 were associated with the endoplasmic reticulum, Golgi, extracellular compartment, mitochondria and nucleus (Figure 10B). F4 was also found to contain more membrane and signal peptide containing proteins than F2; 610 proteins (30.7%) versus 337 proteins (24.5%), and 165 proteins (8.3%) versus 44 proteins (3.2%), respectively (Figure 10C).

GO analysis of the proteins in fractions F2 and F4 also showed differences in the Biological Process GO terms enriched. The top three GO terms for F2 were ‘nucleotide-sugar biosynthetic process’ (GO:0009226), ‘actin polymerization or depolymerization’ (GO: 0008154), and ‘regulation of actin cytoskeleton organization’ (GO:0032956) (Figure 10D). For F4, the top three GO terms were ‘peroxisome organization’ (GO:0007031), ‘protein targeting to ER’ (GO:0045047), and ‘establishment of protein localization to ER’ (GO:0072599) (Figure 10E).

## Discussion

In this study, we developed a method for isolating intracellular vesicles from the late blight pathogen *P. infestans* and identified their associated proteins by mass spectrometry. We employed cushioned bottom-loaded gradient ultracentrifugation to successfully isolate intracellular vesicles, based on the observations that i) proteins migrating to the buoyant fractions F2, F3 and F4 coincided with the presence of vesicles; ii) the virtual absence of vesicles in the dense fraction F6; and iii) the absence of both protein migration and vesicle presence on pretreatment of the buoyant fractions with the detergent Triton X-100, coinciding with increased protein accumulation in the denser fractions, including F6. Together, these observations suggest that intracellular vesicles migrate up the density gradient to fractions of lower density, separated from free proteins/protein aggregates and cellular debris, which accumulate at the bottom of the gradient in higher density fractions.

Mass spectrometry revealed that the presence of homologues of well-characterised membrane and vesicle marker proteins (e.g., TETs, Rabs, SNAREs) commonly associated with EVs were enriched within the buoyant fractions. This suggests that these vesicles have been captured during their biogenesis and trafficking within the cell. In addition to the membrane-associated proteins, the secreted effectors PITG_04314 and PITG_01029 were also identified in the proteomics dataset, indicating that soluble cargo proteins were also isolated using this method. The robust dataset presented here provides a valuable resource that can be mined for potential membrane and cargo markers from *P. infestans* intracellular vesicles. Additionally, our method can be adapted for future studies to look at vesicles with specific buoyant characteristics in detail.

In this study, we focused analysis on proteins commonly enriched in all three buoyant fractions over the denser fraction F6. Rab proteins are well known for their role in intracellular vesicle trafficking ^36,37^, with their presence here suggesting enrichment of diverse intracellular vesicles: Rabs associated with ER-Golgi transport (Rab 1, 2, 18), trans-Golgi network (TGN)-plasma membrane transport (Rab 6, 8, 11), early endosomes mediating clathrin (Rab5) and caveolin (Rab21) pathways ^38^, and late endosomes/lysosome/phagosome (Rab7, Rab 32) were all identified in the buoyant fractions.

Rabs can act as markers for recognition of target membranes during vesicle docking and membrane tethering prior to the initiation of fusion. This recognition is mediated by tethering factors, such as the Golgi-localised complex Vps52/53/54 (homologues of all three of which were identified here) which recognises Ypt6p, the yeast homologue of Rab6. This recognition is required for the initial tethering of vesicles to the trans-Golgi network during retrograde trafficking, bringing the membranes into close proximity for subsequent fusion mediated by SNAREs ^39–41^.

SNAREs mediate the majority of membrane fusion events during exocytosis and endocytosis; they are characterised by 60-70aa SNARE motifs, of which there are four types, Qa, Qb, Qc and R, distributed between vesicle and target membranes, bringing the opposing membranes together on interaction to form the SNARE complex ^41^.

Interestingly, the protein content of the three buoyant fractions were not completely overlapping; this suggests the method used here can separate distinct vesicle populations, as also indicated by the presence of numerous Rabs associated with diverse intracellular vesicle populations. In support of this, comparison between buoyant fractions F2 and F4, representing the extremes of the buoyant fractions analysed in this study, showed considerable differences in the proteins associated with them. Of particular interest are the potential vesicle membrane markers identified amongst these proteins. Present in both fractions were three of the five *P. infestans* tetraspanin homologs; PiTET1 (PITG_03954), PiTET2 (PITG_03950) and PiTET3 (PITG_12224), with PiTET3 significantly more abundant in fractions F2. Similarly, two transmembrane MARVEL domain proteins (PiMDP1 and PiMDP2; PITG_13660 and PITG_13661, respectively) previously identified in *P. infestans* extracellular vesicles ^16^ were also found in fractions F2 and F4, with PiMDP2 enriched in fraction F2.

We propose collection and analysis of more refined fractions over a larger portion of the gradient would reveal the densities at which the PiTET1, PiTET2 and PiMDP1 accumulate. Although this was beyond the scope of the study, the present data demonstrates how this method could be applied to separate distinct vesicle populations associated with specific membrane markers and cargoes. This would aid in the better understanding of the process of protein secretion in *P. infestans* and the development of novel control methods aimed at selectively disrupting this process.

Looking forward, epitope tagging of potential vesicle membrane markers identified in this study would enable further enrichment and purification. Immunoprecipitation of the gradient fractions in which they are most abundant would enable interrogation of protein cargo. Pairing this with markers for specific intracellular localisations, such as the Rab proteins, presents the possibility of identifying the intracellular pathways through which proteins are packaged, trafficked and secreted.

Combining this work with *Phytophthora infestans* EV proteomics will be a powerful tool to further unravel how effector proteins are sorted, transported and secreted. Improving our understanding of how *Phytophthora infestans* infects the potato host will reveal new targets against which pathogen management strategies can be developed and tested to curb the devastating crop losses caused by *P. infestans*.

## Acknowledgements

This work was supported by the European Research Council (https://ror.org/0472cxd90, 787764) and the Biotechnology and Biological Sciences Research Council (https://ror.org/00cwqg982, BB/S003096/1).

The authors would like to thank Dr Piers A. Hemsley for advice on proteomics sample processing; the FingerPrints Proteomics Facility at the University of Dundee; Professor Liz Miller and Dr Laura Spinelli for advice on data analysis and comments on the manuscript; Lydia Welsh for technical support; Dr Petra Boevink for assistance during acquisition of funding.

## Author contributions

J.P., P.R.J.B. and S.C.W. conceived of and designed the research; J.P., S.C. and C.H.H. performed and validated the research; J.P. and S.C.W contributed new reagents/analytical tools; J.P. analysed the data and created visualisations; J.P. wrote the paper with input from all co-authors. P.R.J.B. and S.C.W. won funding for the research.

## Supplementary Figures and Table

**Supplementary Figure S1.**
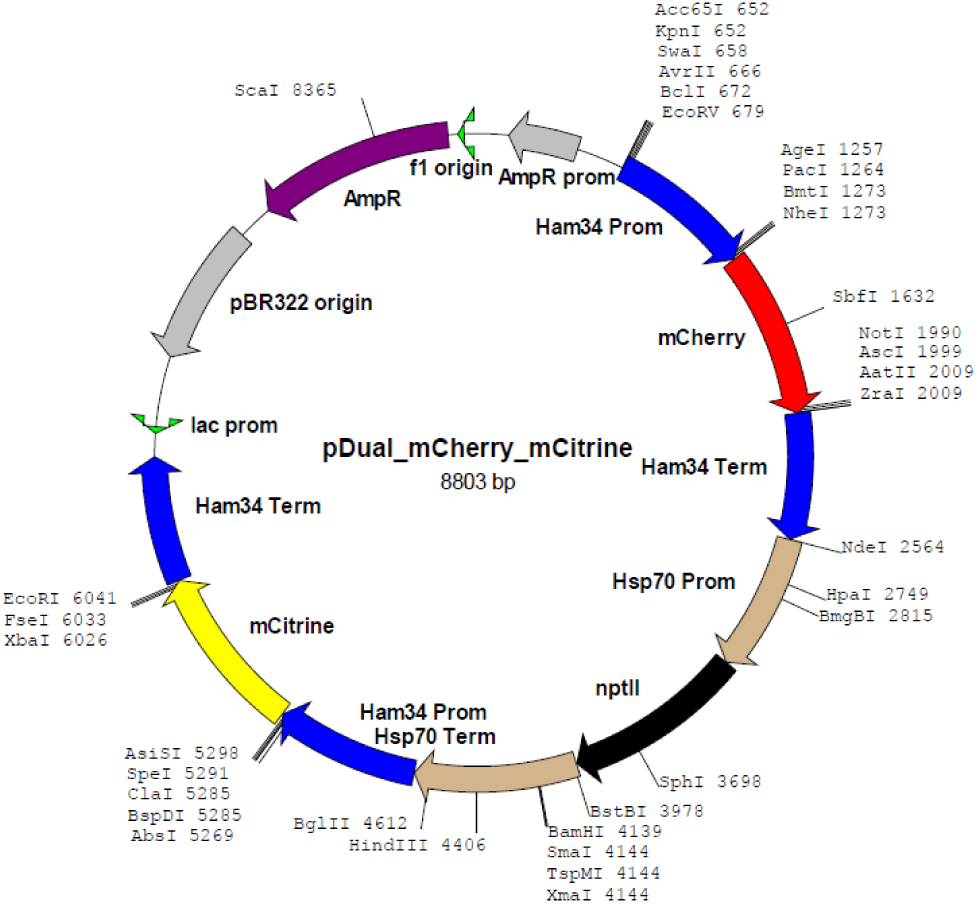
Map of the *P. infestans* expression vector pDual_mCherry_mCitrine constructed for simultaneous expression of mCherry-tagged and mCitrine-tagged proteins.

**Supplementary Figure S2.**
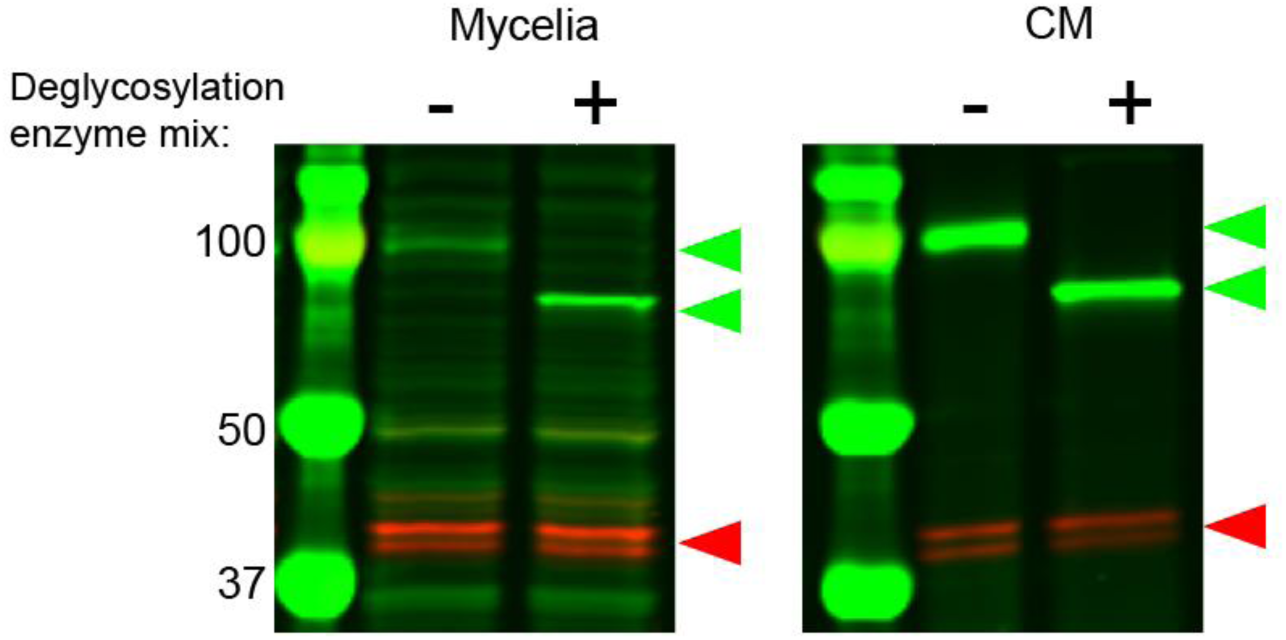
Deglycosylation size shift assay for PITG_04314-mCherry (red arrow) and PITG_01029-mCitrine (green arrows) in (A) *in vitro* grown mycelia and (B) the corresponding conditioned media (CM). “-” = no treatment, “+” = treated with deglycosylation enzyme mix. Size markers (kDa) are shown at left.

**Supplementary Figure S3.**
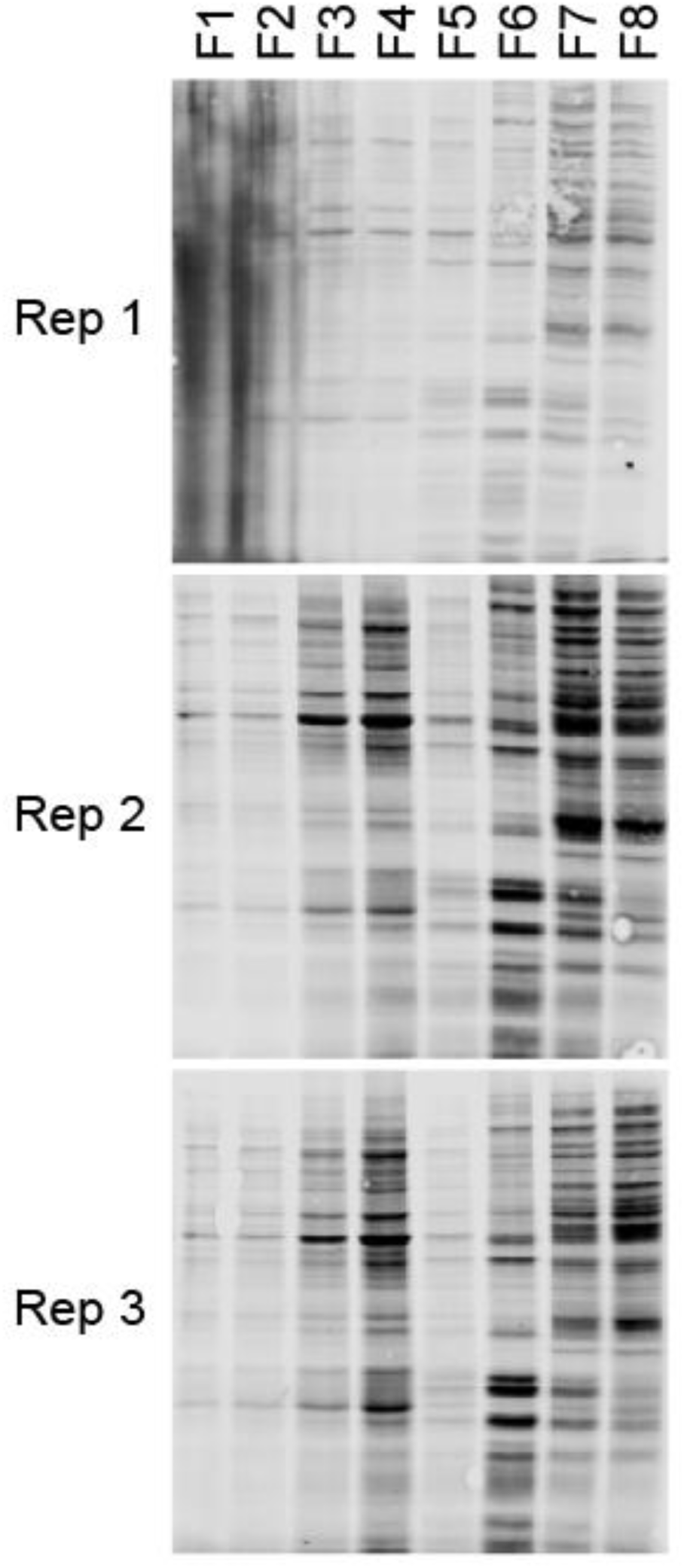
Total protein stain of western blots (shown in Figure 3A) of fractions from three replicate gradients.

**Supplementary Figure S4.**
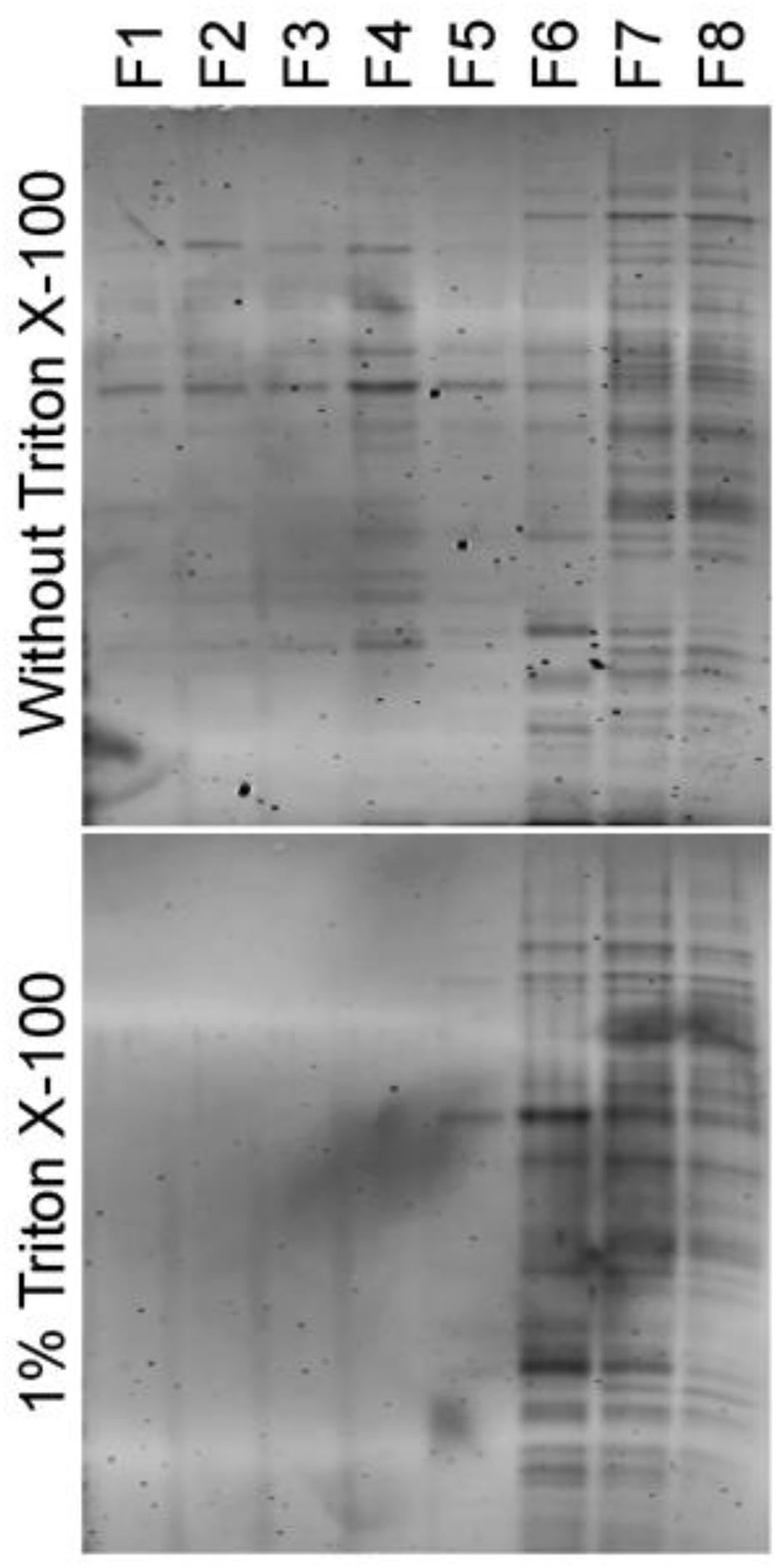
Total protein stain of western blots (shown in Figure 4A and B) of fractions with and without pretreatment with 1% Triton X-100.

**Supplementary Figure S5.**
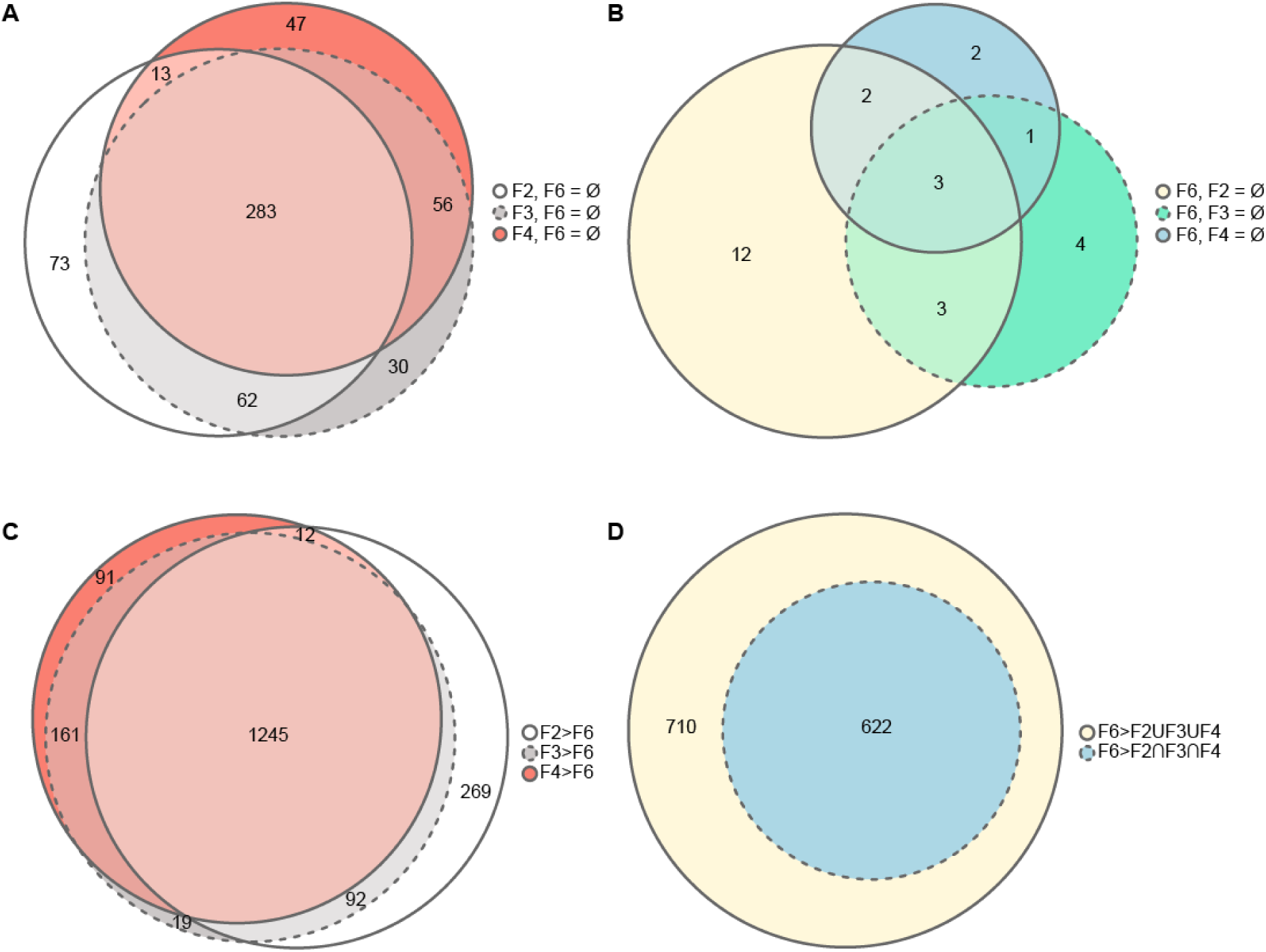
Euler diagrams showing overlap of datasets. (A) Overlap of proteins present (detected in all three biological replicates) in fractions F2, F3 and F4 whilst also absent (not detected in all three biological replicates) in F6 (F6 = Ø). (B) Overlap of proteins present in F6 (detected in all three biological replicates) whilst also absent (not detected in all three biological replicates) in each buoyant fraction (F2 = Ø, 3 F3 = Ø, F4 = Ø). (C) Overlap of proteins in fractions F2, F3 and F4 where Log2FC over F6 was greater than or equal to 0.6 and deemed statistically significant (tested by One-way ANOVA with Tukey’s HSD test (FDR <0.05 and q-value <0.05)). (D) Overlap of proteins in fraction F6 where Log2FC of proteins found in at least one of the buoyant fractions (F2ՍF3ՍF4) over F6 and where Log2FC of proteins found in all buoyant fractions (F2ՈF3ՈF4) was less than or equal to -0.6. Statistical significance tested by One-way ANOVA with Tukey’s HSD test (FDR <0.05 and q-value <0.05).

**Supplementary Figure S6.**
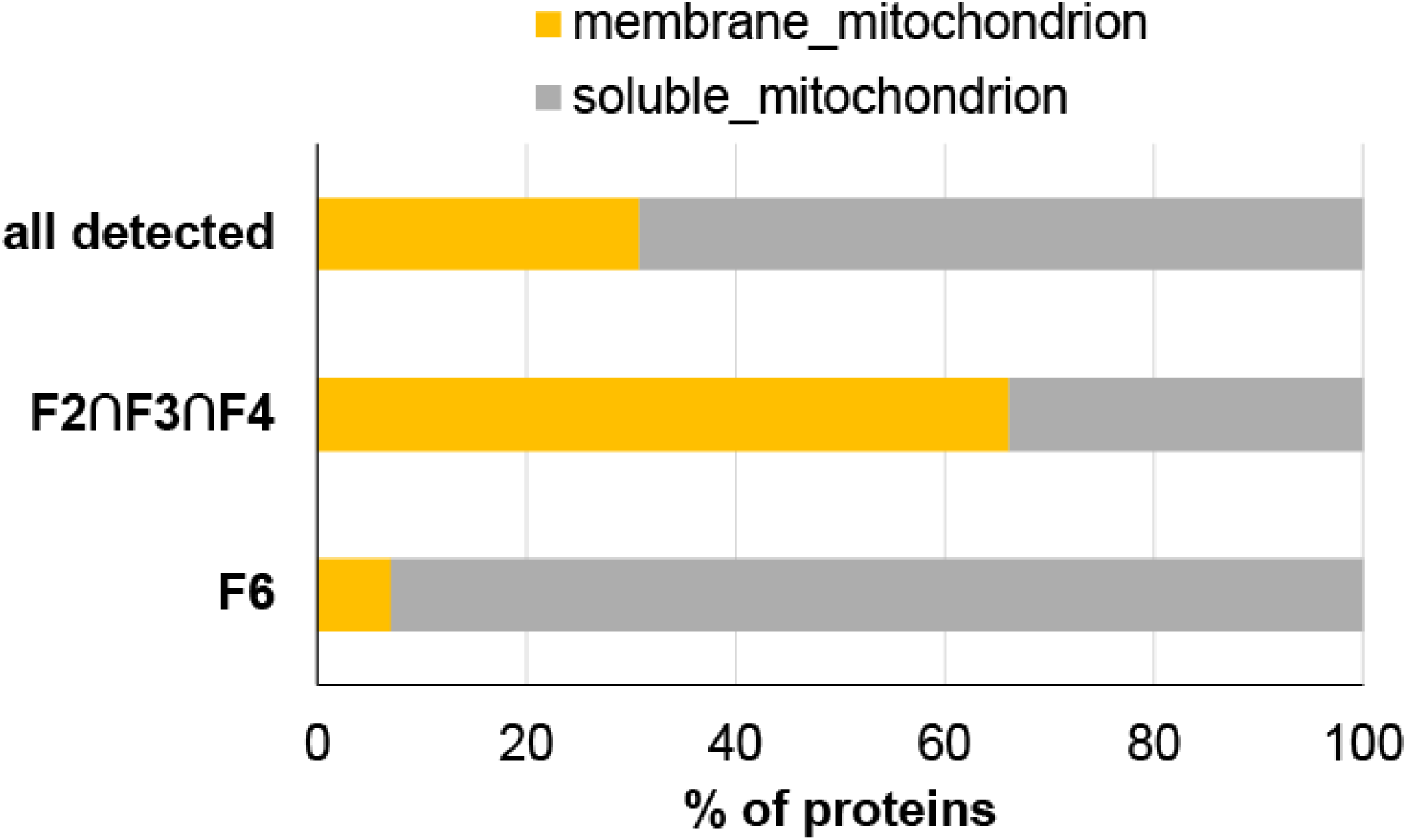
Percentage of predicted mitochondrial membrane and soluble proteins found in the background set of all detected proteins (1,072 total), enriched in buoyant fractions (‘F2∩F3∩F4’) (319 total), and enriched in F6 (71 total).

**Supplementary Table 1.**
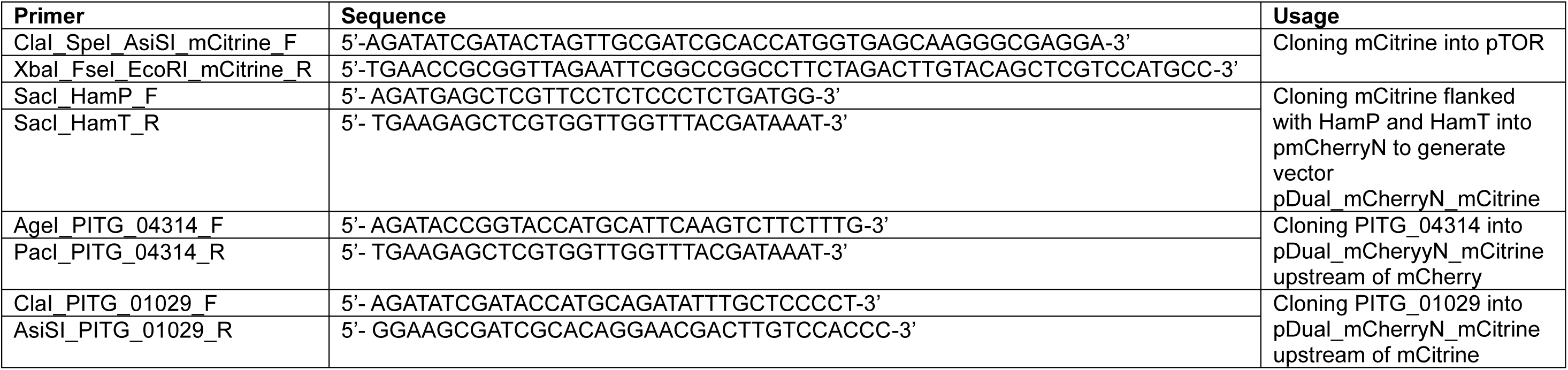
Sequences of primers used to construct vector pDual_mCherryN_mCitrine and to clone PITG_04314 and PITG_01029 into the vector for dual expression in *P. infestans*.

## Supplementary Datasets

**Supplementary Dataset 1**

Tab A: Entire dataset of all proteins detected in the study (6,685 protein groups in total).

Tab B: High confidence proteins commonly enriched within the buoyant fractions F2, F3 and F4 (1,528 protein groups).

Tab C: High confidence proteins enriched within the denser fraction F6 (625 protein groups).

**Supplementary Dataset 2**

Tab A: List of 2,335 signal peptide containing proteins combined from FungiDB, Rafaelle *et al*. ^31^ and Meijer *et al*. ^30^ against which the dataset from the present study was compared against.

Tab B: List of signal peptide containing proteins (+SP) identified within the total background set of 6,685 protein groups. Those enriched within the buoyant fractions and within fraction F6 are indicated in columns ‘enriched_buoyant’ and ‘enriched_F6’, respectively.

Tab C: List of verified secreted proteins published in Meijer *et al*. ^30^ identified within the total background set of 6,685 protein groups. Those enriched within the buoyant fractions and within fraction F6 are indicated in columns ‘enriched_buoyant’ and ‘enriched_F6’, respectively.

**Supplementary Dataset 3**

Tab A: ShinyGO output for Biological Processes enriched within the buoyant fractions.

Tab B: Collapsed list of proteins within top three Biological Processes GO terms returned from ShinyGO for buoyant fractions with InterPro and Pfam domain descriptions.

**Supplementary Dataset 4**

Tab A: ShinyGO output for Biological Processes enriched within F6.

Tab B: Collapsed list of proteins within top three Biological Processes GO terms returned from ShinyGO for F6 with InterPro and Pfam domain descriptions.

**Supplementary Dataset 5**

Tab A: List of 1330 protein groups differentially more abundant in F2.

Tab B: List of 1909 protein groups differentially more abundant in F2.

Tab C: List of 44 proteins only present in F2.

Tab D: List of 75 proteins only present in F4.

Tab E: ShinyGO output for Biological Processes enriched within F2.

Tab F: ShinyGO output for Biological Processes enriched within F4.

